# Structural elements facilitate extreme long-range gene regulation at a human disease locus

**DOI:** 10.1101/2022.10.20.513057

**Authors:** Liang-Fu Chen, Hannah Katherine Long, Minhee Park, Tomek Swigut, Alistair Nicol Boettiger, Joanna Wysocka

## Abstract

Enhancer clusters overlapping disease-associated mutations in Pierre Robin sequence (PRS) patients regulate *SOX9* expression at genomic distances over 1.25 megabases. We applied optical reconstruction of chromatin architecture (ORCA) imaging to trace 3D locus topology during PRS-enhancer activation. While we observed pronounced changes in locus topology between cell-types, analysis of single chromatin fiber traces revealed that these ensemble-average differences arise not from the presence of cell-type unique conformations, but through changes in frequency of commonly sampled topologies. We further identified two CTCF-bound elements, internal to the *SOX9* topologically associating domain, which are positioned near its 3D geometric center and bridge enhancer-promoter contacts in a series of chromatin loops. Ablation of these elements results in diminished *SOX9* expression and altered domain-wide contacts. Polymer models with uniform loading across the domain and frequent cohesin collisions recapitulate this multiloop, centrally clustered geometry, suggesting a mechanism for gene regulation over ultralong ranges.

**Four short bullet points that convey the key message of the paper:** *SOX9* domain topology dynamically changes during a developmental transition

Structural elements promote TAD-wide interactions, stripe formation and transcription

Structural elements are CTCF-dependent and situated centrally in the 3D TAD structure

Polymer simulations of multi-loop model best recapitulate topological features

## Introduction

During development, genes are activated in a dynamic manner as part of a regulatory cascade that integrates responses to the external cellular signaling environment and cell-intrinsic lineage context through the combinatorial action of transcription factors (TFs). Much of this integration occurs at non-coding gene regulatory elements called enhancers, which are short DNA sequences encoding for clusters of TF binding sites that act to drive transcription of their target genes, often across large genomic distances (**Furlong & Levine, 2018**; **Long et al., 2020**). The mechanisms by which enhancers regulate gene expression, especially across vast genomic distances, are an active area of inquiry, with much recent attention focused on the role of topologically associating domains (TADs). TADs were originally described from chromosome conformation capture based studies as self-interacting domains that are enriched for interactions within compared to between domains (**Dixon et al., 2012**; **Nora et al., 2012**). More fine-scale mapping of TADs has uncovered increasing details of architecture within these domains, including contact points (also known as off-diagonal dots or loops), architectural stripes (also known as flares or lines) and sub-TADs (**Hsieh et al., 2020**; **Krietenstein et al., 2020**). Functionally, TADs appear to restrict enhancer activity within a given locus. For example, regulatory reporter sensors can capture enhancer activity when inserted within the same TAD as strong enhancers, but not when the reporter insertion lies outside of the domain (**Anderson et al., 2014**; **Symmons et al., 2014**). Furthermore, disruption of TAD organization, through large genomic deletions and inversions involving TAD boundaries, has been implicated in mis-regulation of gene expression and human disease (**Bruijn et al., 2020**; **Franke et al., 2016**; **Laugsch et al., 2019**; **Lupiáñez et al., 2015**).

The topological properties of TADs are thought to arise from the active process of loop extrusion by the cohesin complex which actively spools DNA to create an extending DNA loop (**Fudenberg et al., 2016**; **Sanborn et al., 2015**). The boundaries of TADs are frequently delimited by the genome-organizing factor CTCF bound in a convergent orientation, which can stall and stabilize cohesin (**de Wit et al., 2015**; **Guo et al., 2015**; **Nora et al., 2020**; **Wutz et al., 2020**). Inducible degradation of cohesin or CTCF leads to the dissolution of TAD structures, though surprisingly, the impact on transcription is only moderate (**Nora et al., 2017**; **Rao et al., 2017**). In addition to its function at boundary elements, CTCF has been shown to play structural and/or gene regulatory roles in other contexts, for example through the formation of regulatory neighborhoods within TADs (**Dowen et al., 2014**) or by mediating enhancer-promoter contacts (**Kubo et al., 2021**; **Schuijers et al., 2018**). Notably, the strongest effects of CTCF-loss on gene expression have been observed for genes associated with promoter-proximal CTCF sites, suggesting a broad role for such sites in regulating transcription (**Kubo et al., 2021**; **Nora et al., 2017**).

Recent work has shown that TADs are dynamic structures that appear as an emergent property across a population of cells. Several FISH-based imaging studies that mapped chromosomal folding at the single-cell, single chromatin fiber level revealed considerable structural heterogeneity of chromatin organization at individual loci (**Bintu et al., 2018**; **Cheng et al., 2020**; **Finn et al., 2019**; **Mateo et al., 2019**; **Takei et al., 2021**). Such structural heterogeneity has also been detected with other methods, such as single cell Hi-C experiments (**Nagano et al., 2013**) and genome architecture mapping (GAM) (**Welch et al., 2020**). Thus, topological organization detected at the population level appears to be a product of variable and dynamically changing structures, with barriers to the active process of loop extrusion favoring certain conformations which consequently become enriched in ensemble-average maps.

The *SOX9* locus provides a biologically interesting and disease-relevant model for studies of long-range gene regulation and 3D genome organization. The *SOX9* gene – which encodes for a developmentally-important TF – is the only protein-coding gene immersed in this enormous gene desert within a 2 megabase (Mb) TAD, one of the largest in the human genome. Distinct non-coding mutations within this domain have been associated with several human congenital diseases affecting development of face, limbs or sex determination, consistent with these tissues being the most sensitive to changes in the SOX9 dosage (**Benko et al., 2009**; **Croft et al., 2018**; **Gonen et al., 2017**; **Kurth et al., 2009**; **Yao et al., 2015**). We previously characterized two extreme long-range enhancer clusters (ECs) at the *SOX9* locus, which we termed EC1.45 and EC1.25 to reflect their distance in megabases to the *SOX9* promoter (1.45 Mb and 1.25 Mb to the *SOX9* promoter, respectively) (**Long et al., 2020**). These ECs overlap with patient mutations associated with the isolated craniofacial disorder called Pierre Robin Sequence (PRS) (**Benko et al., 2009**; **Logjes et al., 2018**) and regulate *SOX9* expression specifically in cranial neural crest cells (CNCCs), a transient embryonic population that gives rise to the majority of craniofacial structures during development (**Bronner & LeDouarin, 2012**; **Long et al., 2020**). Interestingly, based on the CNCC-specific enrichment of active enhancer chromatin marks, our previous work nominated a third candidate regulatory element overlapping PRS patient deletions (termed E1.35), located 1.35Mb from the promoter between EC1.45 and EC1.25 (**Long et al., 2020**). In contrast to EC1.45 and EC1.25, however, this element lacked autonomous enhancer activity in reporter assays in human CNCCs or mice, leaving its potential function in *SOX9* regulation unresolved. Furthermore, it remains unclear how EC1.45 and EC1.25, which represent some of the longest-range regulatory elements functionally characterized in the human genome, can achieve precise *SOX9* regulation across genomic distances of over 1.2 Mb. Such precision is indeed required for normal craniofacial development, as we demonstrated that even small, 10-15% changes in *SOX9* dosage in the developing face are sufficient to result in subtle, but reproducible changes in lower jaw morphology (**Long et al., 2020**).

Here, we use optical reconstruction of chromatin architecture (ORCA) (**Mateo et al., 2019**) to investigate the topological folding of the *SOX9* locus in two cellular states: (i) human embryonic stem cells (hESCs), where ECs are inactive and *SOX9* is lowly transcribed, and (ii) CNCCs, where both ECs and *SOX9* are highly active. At the ensemble level, we observe pronounced changes in locus topology between the two states. Specifically, sub-TAD structures and a paucity of long-range interactions with the distal region of the TAD are seen in hESCs, whereas nearly uniform interactions throughout the entire TAD and appearance of two architectural stripes are seen in CNCCs. Analysis of our ORCA data at the single chromatin fiber level reveals that these differences arise through cell-type specific changes in frequency of topologies commonly sampled by both cell-types. We further show that the two architectural stripes observed in CNCCs form at TAD-internal regions corresponding to an E1.35 and *SOX9* promoter-proximal CTCF site, which we refer to as ‘stripe-associated structural elements’ (SSEs). These SSEs regulate *SOX9* expression in a CTCF-dependent manner. However, rather than selectively promoting interactions between specific enhancer-promoter pairs, the two structural elements broadly facilitate contacts throughout the entire TAD, leading to increased frequency of multiway interactions between enhancers at the locus and *SOX9* promoter. We demonstrate that the structural elements are centrally positioned within the 3D topology of the domain, suggesting a plausible explanation for their broad role in promoting intra-TAD contacts. Polymer simulations of loop extrusion show that locus topologies and central positioning of the structural elements are better approximated by a ‘multi-loop’ model, as compared to the previously proposed ‘reel-in’ model of architectural stripe formation (**Vian et al., 2018**). Together, utilizing single-trace imaging data, we uncover structural elements at a human disease locus that facilitate domain-wide interactions, reside centrally within the domain, and are required for robust regulation of gene expression. Furthermore, leveraging polymer simulations we propose a multi-loop model of locus folding that explains observed topologies, stripe formation and central structural element positioning.

## Results

### Pronounced changes in *SOX9* locus topology accompany neural crest cell differentiation

To investigate molecular mechanisms governing human craniofacial development, facial progenitor cells called cranial neural crest cells (CNCCs) can be derived *in vitro* from human embryonic stem cells (hESCs) (**Bajpai et al., 2010**; **Prescott et al., 2015**). During this differentiation, the expression of an important developmental transcription factor *SOX9* is upregulated around 6-fold in CNCCs compared to hESCs (**Figure 1A-B**). Multiple putative enhancers are activated across the 2 Mb gene desert surrounding the *SOX9* gene during this transition, including two extremely long-range enhancer clusters (EC1.45 and EC1.25) that map within the PRS disease locus and regulate *SOX9* expression in CNCCs (**Figure 1C and Supplementary Figure 1A**) (**Long et al., 2020**). Given the large size of the *SOX9* topologically associated domain (TAD) and the vast distances across which the PRS ECs function, we sought to leverage this locus to explore the mechanisms by which extreme- distal enhancers function in 3D space.

**Figure 1.**
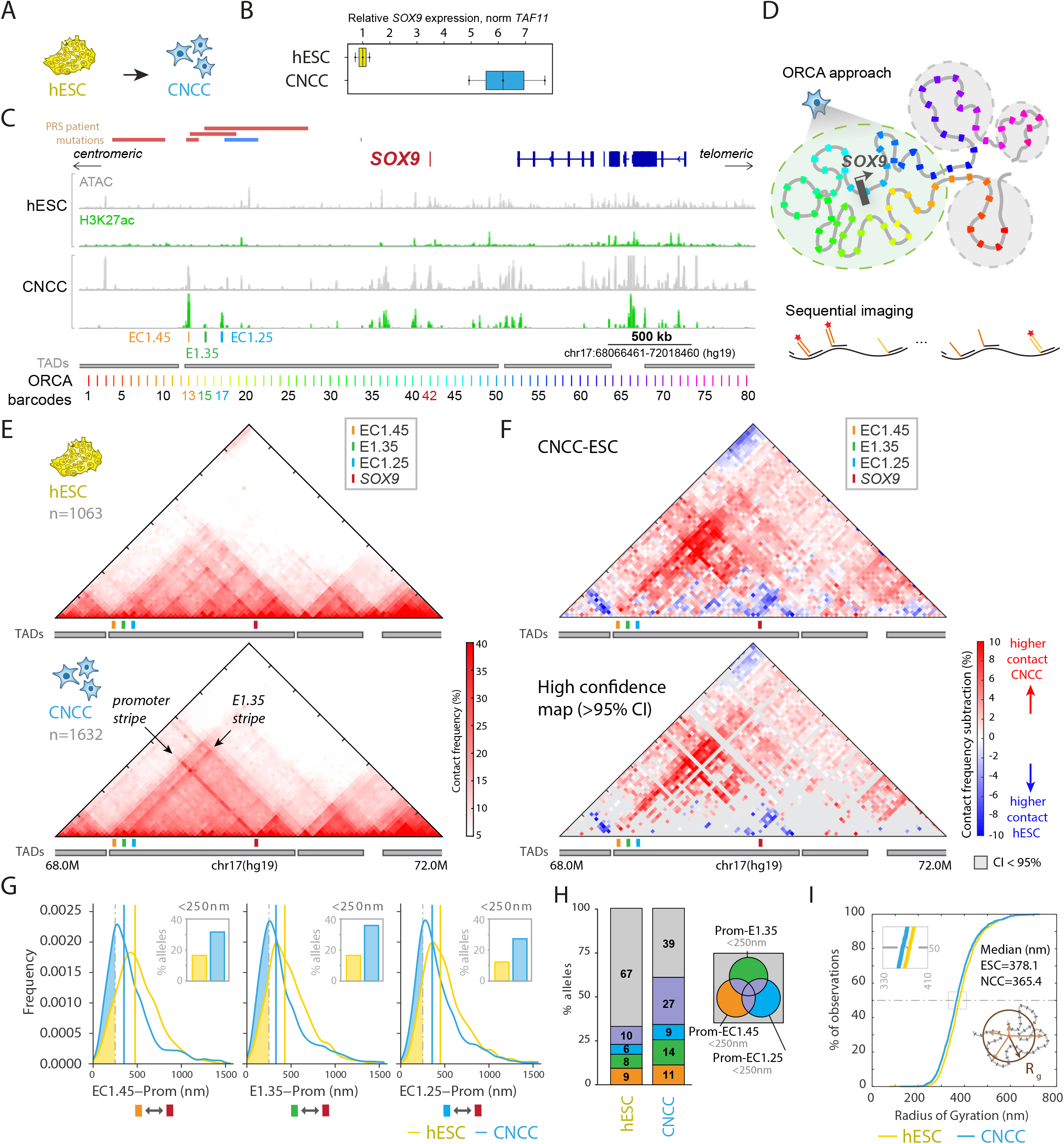
ORCA imaging reveals changes to domain topology during cranial neural crest cell differentiation. A) Schematic of hESC to CNCC differentiation. B) Boxplot of *SOX9* expression measured by RT-ddPCR normalized to *TAF11* with hESC median set to 1. n=3. C) ATAC-seq and H3K27ac ChIP-seq data from hESCs and CNCCs for a 4 Mb locus surrounding the *SOX9* gene. Enhancer clusters EC1.45 and EC1.25 are highlighted along with the nearby E1.35 element. 80 ORCA barcodes are annotated across the domain. PRS patient mutations are highlighted including large deletions (red) and translocation breakpoint cluster (blue) (**Amarillo et al., 2013**; **Benko et al., 2009**; **Gordon et al., 2014**); TADs are indicated in grey. ORCA barcodes: chr17:68066461-72018460 (hg19). D) Schematic of ORCA strategy. Barcodes are imaged sequentially. E) Contact frequency maps for pairwise distances <250 nm from ORCA imaging data for hESC (upper) and CNCCs (lower). Arrows highlight two domain-spanning stripes in CNCCs. F) Contact frequency subtraction maps for CNCC-hESC (upper) and high confidence map highlighting robust significant changes, pixels are shown with confidence interval > 95% (lower). G) Frequency plots showing pairwise distance distributions for hESC (yellow) and CNCC (blue) for EC1.45, E1.35 and EC1.25 to the *SOX9* promoter from left to right. Percentage of alleles with respective pairwise distances <250 nm are shown as an inset. H) Percentage of alleles with promoter proximity to EC1.45, E1.35 and/or EC1.25 including multiway interactions for hESCs and CNCCs. I) Cumulative distribution function plot of the radius of gyration for the *SOX9* TAD for hESC (yellow) and CNCCs (blue).

We first aimed to visualize chromosomal folding of the *SOX9* locus in both hESCs and CNCCs. ‘Optical reconstruction of chromatin architecture’ (ORCA) is a multiplexed fluorescence *in situ* hybridization (FISH) methodology that utilizes sequential locus imaging to reconstruct chromosome topology for a region of interest (**Figure 1D**) (**Mateo et al., 2019**). We designed ORCA probes to label 80 5kb-regions at 50kb intervals across a 4 Mb genomic region encompassing the *SOX9* regulatory domain and flanking TADs (**Figure 1C-D**, chr17:68,066,461-72,018,460, hg19). Importantly, we designed our set of probes to overlap with the *SOX9* promoter, enhancer clusters EC1.45 and EC1.25, as well as the enhancer-associated element E1.35 (**Long et al., 2020**) (**Figure 1C**).

We imaged ∼4 Mb of chromosome topology for 100s of ORCA traces for both hESCs and CNCCs. Pairwise distances between all 80 regions across the domain were calculated from x-y-z positions for each probe set. To examine the population average topology, the percentage of pairwise interprobe distances below 250 nm were plotted as a ‘contact frequency’ map (**Figure 1E**; see also **Supplementary Figure 1B** for absolute average pairwise distances). In hESCs, regions of self-interacting chromatin matching TAD structures identified from Hi-C assays were clearly observed, including the large (∼2 Mb) TAD harboring *SOX9* (**Akgol Oksuz et al., 2021**) (**Figure 1E**, upper and **Supplementary Figure 1D**). In agreement with our previously published Capture-C data from the *SOX9* promoter (**Long et al., 2020**), we observed that in hESCs the *SOX9* promoter preferentially interacts with the telomeric side of the TAD, with fewer long-range interactions across the remainder of the domain (**Figure 1E**, upper). Within the nearly 2 Mb *SOX9* TAD, smaller blocks of interacting chromatin are also observed in hESCs, reminiscent of sub-TAD structures (**Akgol Oksuz et al., 2021**; **Rao et al., 2014**). These patterns emerge from the population average structure and may reflect a preferential folding of the locus into smaller blocks of self-interacting chromatin in hESCs within the overarching constraints of the larger TAD. In contrast to hESCs, ORCA contact frequency maps in CNCCs revealed nearly uniform levels of interactions across the *SOX9* TAD, with two striking stripe-like features emanating from the ORCA probe set overlapping the *SOX9* promoter and the E1.35 region (**Figure 1E**, lower – see arrows).

To compare the average structures between hESCs and CNCCs, we generated subtraction maps for the contact frequency and absolute distance maps (**Figure 1F**, upper and **Supplementary Figure 1C**). These maps highlight that shorter-range interactions around the *SOX9* promoter are more common in hESCs (closer to the horizontal line), while longer-range interactions become more prevalent in CNCCs. Subtraction maps further highlight the two stripe features in CNCCs, and also reveal an increased interaction along the centromeric TAD boundary in CNCCs. Taking advantage of the single-trace nature of the ORCA data, we performed bootstrapping analysis to determine which changes were statistically significant, which further highlighted the switch from short-range interactions on the telomeric side of the *SOX9* TAD in hESCs, to long-range TAD-spanning interactions in CNCCs (**Figure 1F**, lower, high confidence map showing pixels with confidence interval > 95% by bootstrapping).

To quantify the increase in long-range interactions observed in CNCCs, we focused on interactions of the *SOX9* promoter region with the EC1.45, E1.35 and EC1.25 regions at the PRS locus. In keeping with the increase in long-range interactions observed in CNCCs, the median distance between the PRS ECs and the *SOX9* promoter was significantly decreased (**Figure 1G**, p< 0.001, K-S test). Furthermore, the frequency of promoter proximity <250 nm to the EC1.45, E1.35 and EC1.25 elements was increased around 2-fold in CNCCs to 31.2%, 35.9% and 27.3%, respectively (**Figure 1G**, insets). When considering all promoter-EC interactions, the proportion of ORCA traces for which the promoter is proximal to any of the three PRS locus elements rises to 60.9% in CNCCs, from 33.0% in hESCs. The majority of the increase is accounted for by an increase in multi-way interactions involving more than one of the PRS locus elements (**Figure 1H**). Together, this suggests that in CNCCs the *SOX9* promoter region frequently comes into close proximity with ECs across extreme long-distances, perhaps forming a hub-like conformation involving simultaneous contacts with multiple distal elements (**Supplementary Figure 1E**, compare right to left).

To determine whether the observed decreases in pairwise distances across the *SOX9* TAD translate to a smaller volume for the domain, we calculated the radius of gyration for hESCs and CNCCs across all observed traces. The radius of gyration is calculated for each allele as a mean of all pairwise distances across the TAD to the calculated center of the domain (**Figure 1I**, lower right inset). This analysis revealed that at the population level, there is a small but significant decrease in the radius of gyration (p < 0.001, Wilcoxon rank sum test) for the *SOX9* domain in CNCCs compared to hESCs (**Figure 1I**). However, this decrease was relatively modest (the overall radius of gyration changed only by around 13 nm) compared to the changes in pairwise distances within the domain, which were often greater than 100 nm (**Figure 1G** and **Supplementary Figure 1C**). Together, our ORCA results reveal dramatic changes in *SOX9* locus topology during neural crest differentiation, with large changes occurring to pairwise interactions within the TAD, while little change was observed to the overall volume of the domain.

### Locus topologies sampled by individual chromatin fibers are not unique to a given celltype, but certain conformations are more common in hESCs or CNCCs

Our initial analysis of combined population heatmaps of *SOX9* locus topology revealed a distinct locus architecture for hESCs compared to CNCCs (**Figure 1E**). We therefore sought to leverage the single-trace nature of our dataset to distinguish between two plausible hypotheses of how these distinct architectures can arise at the population level: (i) through the presence of chromatin conformations unique to each cell-type, or (ii) through cell-type specific changes in frequency of conformations commonly sampled by both cell-types. We restricted our analysis to the *SOX9* TAD and first focused on all pairwise distances for the 42 barcodes across this region. Using t-distributed stochastic neighbor embedding (tSNE) analysis, we reduced the dimensionality of the pairwise distance matrix to visualize each trace in two dimensions (**Supplementary Figure 2A**). No clear separation or clustering was observed for the two cell states. We reasoned that many regions within the TAD may not exhibit distinctive features that contribute to cell-type specificity of locus structure. We therefore reduced our analysis to all pairwise distances of the *SOX9* promoter to the surrounding TAD, as these interactions represent some of the most dramatic changes, we observed at the population level (**Figure 1F**). Focusing on only the *SOX9* promoter pairwise distances, we observed a slight separation between hESC and CNCC traces on the tSNE plot. However, the overall topological conformations sampled by the promoter in both cell-types appear to be highly overlapping (**Figure 2A**).

**Figure 2.**
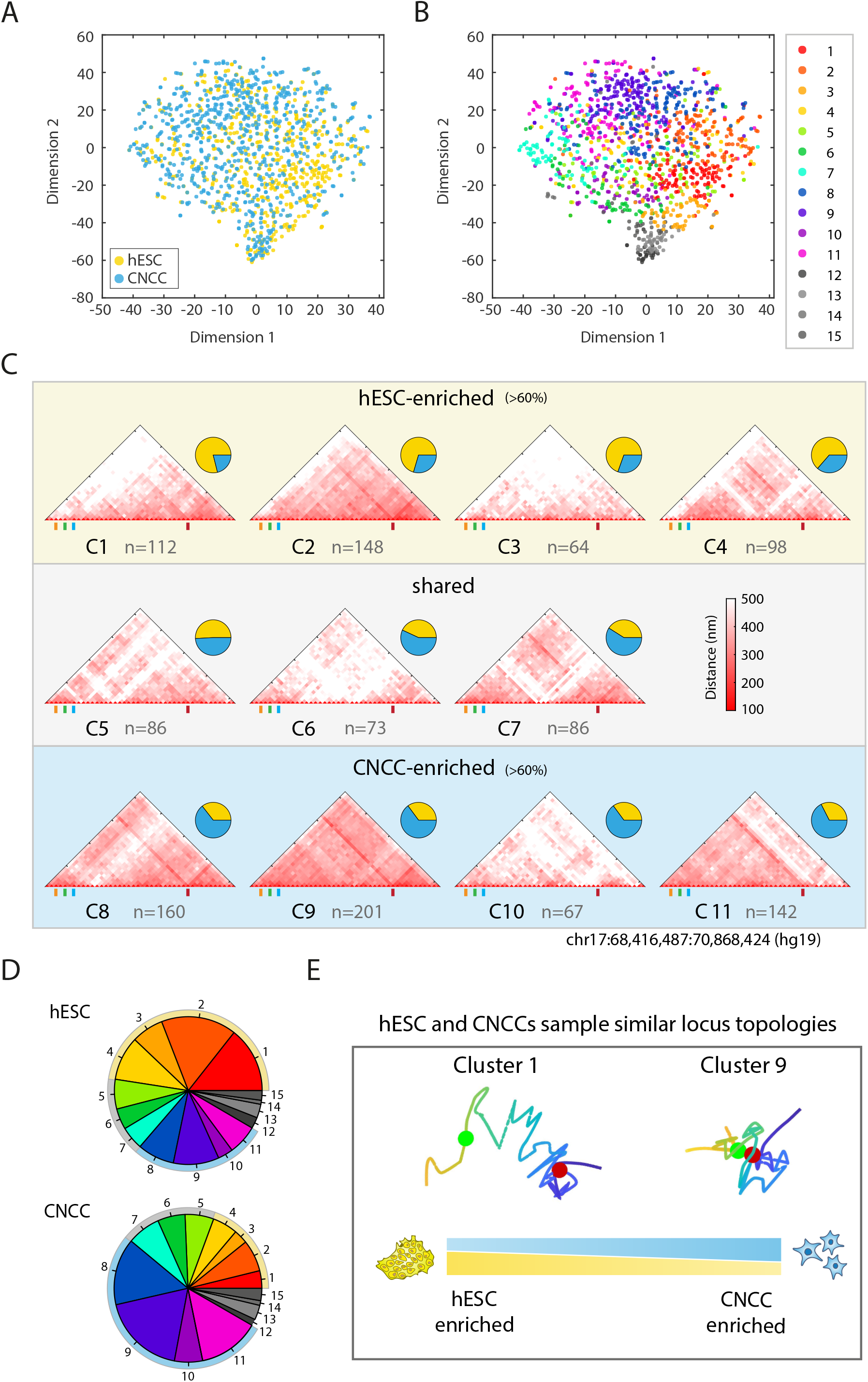
Distinct 3D domain conformations are enriched for, yet shared between, hESCs and CNCCs. A) A tSNE plot representing all pairwise distances of the *SOX9* promoter to the remainder of the TAD (data from Figure 1). B) tSNE plot from Figure 2A indicating 15 clusters from K-means clustering. C) Absolute pairwise distance maps from all traces for each cluster from Figure 2B. Pie charts indicate proportions of traces for each cluster from hESC (yellow) or CNCC (blue). Clusters that are hESC-enriched or CNCC-enriched are highlighted. D) Pie charts showing proportions of traces in each K-means cluster for hESC (upper) and CNCC (lower). E) Schematic illustrating the shared locus topology for hESC and CNCC. Cell-type enrichment for certain structures is highlighted by MDS traces for exemplar hESC or CNCC-enriched structures.

To explore if certain locus conformations are more frequent in one cell-type or the other, we performed k-means clustering of the *SOX9* locus topology. Here, we focused the analysis only on the pairwise interactions of the *SOX9* promoter with other regions within the TAD. Elbow analysis suggested that three clusters may be the optimal number for segregating distinct structural subsets in the combined hESC-CNCC dataset (**Supplementary Figure 2B**). Three clusters were therefore determined from the hESC-CNCC pairwise distance data and visualized on the tSNE plot (**Supplementary Figure 2C**). One of the three clusters (cluster 2) was relatively hESC-enriched (61%) and one (cluster 1) was CNCC-enriched (56%). Plotting the ensemble average heatmaps for each of the three clusters revealed a clear difference in locus architecture between the clusters (**Supplementary Figure 2D**). In particular, cluster 1 showed uniform interactions across the TAD with two clear stripes, reminiscent of the ensemble map from CNCCs, while cluster 2 showed sub-TAD like structures and enrichment of promoter-proximal interactions, reminiscent of the ensemble map from hESCs.

To further sub-stratify distinct classes of structure within the *SOX9* TAD, we allowed for a larger number of clusters (15) and analyzed 11 clusters which had greater than 50 single traces within them (**Figure 2B** and **Supplementary Figure 2E**, clusters 1-11). The 11 clusters appeared to occupy distinct regions of the tSNE map, with 4 clusters enriched in hESC and 4 enriched in CNCC (**Figure 2C-D** and **Supplementary Figure 2E**). The remaining 3 clusters appeared to be present at comparable frequencies in both cell states. hESC-enriched ensemble heatmaps exhibited features reminiscent of the hESC population structure, including distinct sub-domain boundaries, and a gradation of interactions strongest proximal to the promoter that becomes weaker at increasing distances (e.g. clusters 1 and 2). By contrast, the CNCC-enriched clusters exhibited domain-wide interactions with the presence of the *SOX9* promoter stripe, and in some cases, also the E1.35 stripe (e.g. clusters 8 and 9) (**Figure 2C**). Taken together, our results are consistent with a model whereby the *SOX9* promoter samples a similar constellation of locus topologies in hESCs and CNCCs, but certain conformations are more common in one of the cell-types, resulting in substantial cell-type specific differences in ensemble average contact maps (**Figure 2E**).

### The PRS locus E1.35 element regulates *SOX9* expression in the absence of enhancer activity

The most distinctive features of the CNCC contact frequency ensemble maps are two intersecting stripes within the TAD. These stripes occur at the ORCA barcodes which overlap with either the *SOX9* promoter region or the E1.35 regulatory element far upstream of *SOX9* (**Figures 1E and 2C**). In our previous work, we characterized the activity of three CNCC-specific regulatory sequences that are ablated in PRS patients. Enhancer clusters EC1.45 and EC1.25 were highly active in CNCCs by luciferase assays and in mouse craniofacial embryonic development as determined by LacZ reporters (**Long et al., 2020**). In contrast, E1.35 did not appear to have autonomous regulatory capacity by either measure, suggesting that it may not function as an enhancer (**Supplementary Figure 3A-B**). The apparent presence of a stripe feature at E1.35 led us to revisit a role for this region in modulation of *SOX9* expression. We generated two independent cell lines with heterozygous deletion of E1.35 and used our published 10X linked-read sequencing to phase the wildtype and deleted alleles with the two copies of the *SOX9* gene. Using allele-specific ddPCR probes, we examined the allelic expression of *SOX9* for wildtype and E1.35 mutant lines during CNCC differentiation and observed a ∼20% reduction of gene expression upon E1.35 loss (**Figure 3A-B**), an effect comparable to the expression reduction we have previously observed for EC1.25 deletion (**Long et al., 2020**) (**Supplementary Figure 3C**). Therefore, despite its lack of canonical enhancer reporter activity, the E1.35 element contributes to *SOX9* regulation in CNCCs.

**Figure 3.**
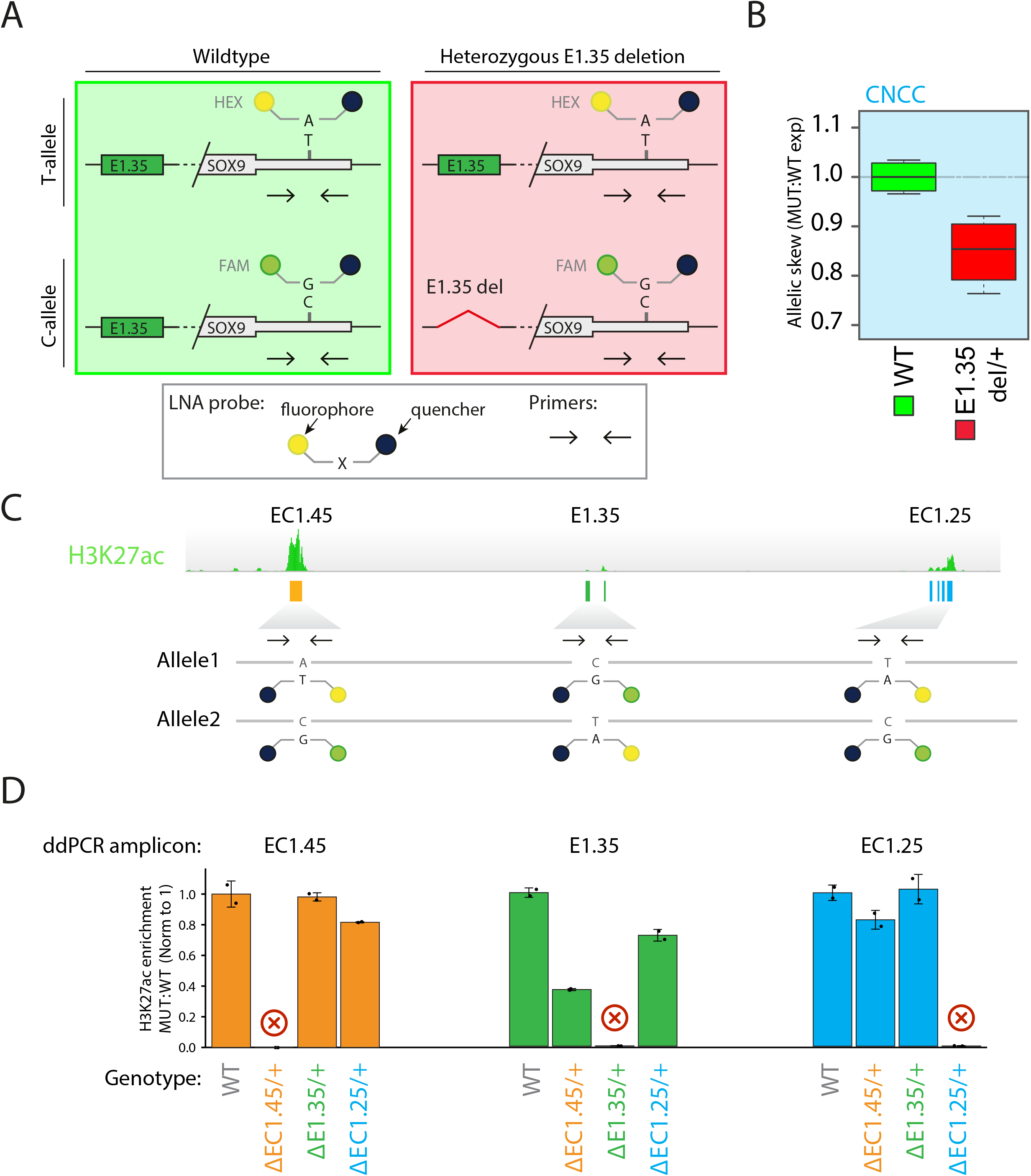
Ablation of E1.35 element affects SOX9 expression, but does not impact the activity of flanking enhancer clusters, as measured by active histone modifications. A) Schematic of allele-specific RT-ddPCR highlighting allele-specific SNPs and LNA probes for wild-type and heterozygous E1.35 deletion cell-lines. B) RT-ddPCR for wild-type (green boxplot) and E1.35 heterozygous deletion (red), plotting mutant to wild-type expression ratio with wild-type normalized to 1. C) Schematic of allele-specific ddPCR for H3K27ac ChIP-ddPCR at EC1.25, E1.35 and EC1.45. D) ChIP-ddPCR for H3K27ac at EC1.45, E1.35 and EC1.25 from wild-type and heterozygous deletions of EC1.45, E1.35 or EC1.25 CNCCs. Ratio of mutant to wild-type ratio signal is plotted, with wild-type normalized to 1.

We hypothesized that E1.35 may contribute to *SOX9* regulation through non-canonical means. Given the genomic location of E1.35 between two strong enhancer clusters, and the presence of chromatin modifications and co-activator binding associated with enhancer activity, we firstly reasoned that E1.35 may act to facilitate or boost EC1.45 and/or EC1.25 activity, similarly to facilitator elements recently described at the alpha globin locus (**Blayney et al., 2022**). To test this hypothesis, we predicted that if E1.35 were acting to boost the activity of the flanking ECs, loss of E1.35 would reduce the activity of EC1.45 and/or EC1.25. We reasoned that levels of H3K27ac could act as a proxy for enhancer activity levels and designed allele-specific probes and ChIP ddPCR primers for each of the three regulatory elements. We performed H3K27ac ChIP-ddPCR in CNCCs that were either wildtype or had a heterozygous deletion for each regulatory region (**Figure 3C**). The resultant ChIP signal was normalized to 1 for wildtype CNCCs for each tested region (EC1.45, E1.35 and EC1.25). Ablation of EC1.45 caused a small reduction in H3K27ac ChIP signal for the EC1.25 region, and a large (∼65%) reduction in signal for the E1.35 element. Similarly, deletion of EC1.25 caused a small reduction in signal at EC1.45 and a slightly larger impact on E1.35 H3K27ac signal (**Figure 3D**). Thus, despite ∼100 kb distance from E1.35 to each flanking EC, deletion of either EC1.45 or EC1.25 results in diminished H3K27ac at E1.35. By contrast, deletion of E1.35 had no impact on H3K27ac signal for either EC1.45 or EC1.25. This result indicates that E1.35 does not act as a booster of PRS locus enhancer activity, at least as measured by their H3K27ac enrichment levels.

### Stripe-associated structural elements (SSEs) at E1.35 and *SOX9* promoter mediate stripe formation and facilitate domain-wide interactions

We next hypothesized that E1.35 may play a structural role to bridge the large genomic distance between the distal region of the TAD and the *SOX9* promoter. In this model, we predicted that E1.35 may influence *SOX9* expression by impacting locus topology, for example by acting as a tethering element between enhancers at the PRS locus and the *SOX9* promoter (**Batut et al., 2022**). Indeed, the TAD-spanning stripe originating from E1.35 in CNCCs may support this theory (**Figure 1E**). Stripe-like patterns (also known as lines or flares) have been observed at many genomic locations by Hi-C-based studies (**Barrington et al., 2019**; **Hsieh et al., 2020**; **Kraft et al., 2019**; **Vian et al., 2018**) and are proposed to form through the process of active loop extrusion stalled by barriers such as CTCF bound in a convergent orientation (**de Wit et al., 2015**; **Fudenberg et al., 2016**; **Guo et al., 2015**). Indeed, utilizing our published CTCF ChIP-seq data, we observed strong CTCF binding at the E1.35 element (**Figure 4A**). Notably, a strong CTCF peak is also observed proximal to the *SOX9* promoter, and the E1.35-associated and promoter-proximal CTCF binding sites are arranged in a convergent orientation (**Figure 4A**). To test if CTCF binding is necessary for the contribution of E1.35 to *SOX9* expression, we generated heterozygous hESC deletion lines for which the CTCF site within E1.35 was ablated; we confirmed loss of CTCF binding in two independent clones by ChIP-qPCR (**Supplementary Figure 4A-B**). Subsequent analysis of *SOX9* allelic expression in CNCCs revealed that loss of the CTCF binding site within E1.35 phenocopies the expression defect seen for the deletion of the entire element, with around 20% reduction (**Figure 4Bi**).

**Figure 4.**
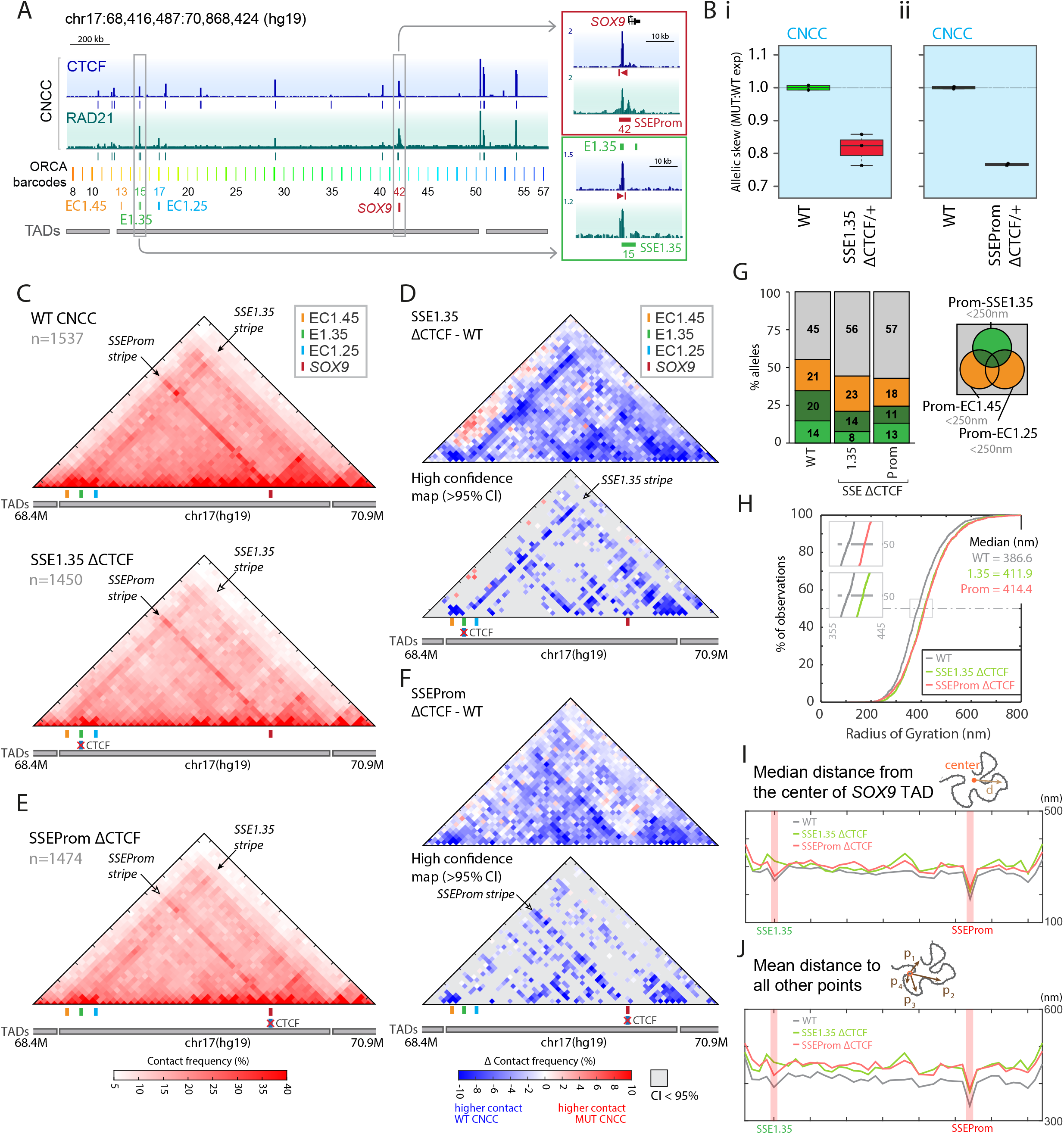
Stripe-associated structural elements are required for TAD-spanning stripes, robust SOX9 expression and central positioning within TAD. A) CTCF and RAD21 ChIP-seq for CNCCs across a 2 Mb locus surrounding the *SOX9* TAD. ORCA barcodes 8-57 are highlighted: chr17:68,416,487:70,868,424 (hg19). B) RT-ddPCR for B) i: E1.35 (SSE1.35) CTCF heterozygous deletion and ii: SSEProm CTCF heterozygous deletion (red box plots), compared to wild-type (WT, green boxplot), plotting mutant to wild-type expression ratio, with wild-type normalized to 1. C) Contact frequency maps for pairwise distances <250 nm from ORCA imaging data for wild-type (upper) and E1.35 CTCF deletion (SSE1.35 Δ CTCF) CNCCs (lower). D) Upper, contact frequency subtraction maps for E1.35 CTCF deletion (SSE1.35 Δ CTCF) compared to wildtype. Lower, high confidence map highlighting robust significant changes, pixels are shown for confidence interval > 95%. E) As C, for SSEProm CTCF deletion CNCCs (SSEProm Δ CTCF). F) As D, for SSEProm Δ CTCF CNCCs. G) Percentage of alleles with promoter proximity to EC1.45, SSE1.35 and/or EC1.25 including multiway interactions for wild-type, SSE1.35 Δ CTCF and SSEProm Δ CTCF CNCCs. H) Cumulative distribution function plot of the radius of gyration for the *SOX9* TAD in wild-type, SSE1.35 Δ CTCF and SSEProm Δ CTCF CNCCs. I) Median distance from the center of the TAD to each point across the domain, averaged across all traces and plotted for each position across the locus. J) The mean distance from a given point to all other points across the domain is plotted for each position across the locus.

To interpret the impact of E1.35 CTCF site ablation on *SOX9* gene expression, we repeated single-trace ORCA imaging of the *SOX9* locus in CNCCs for both wildtype and E1.35 CTCF homozygous knock-out lines (E1.35 ΔCTCF). Contact frequency maps revealed complete loss of the PRS locus-associated TAD-spanning stripe, and an overall reduction in contact frequency across the domain (**Figure 4C**, lower; see lighter red across the domain for the E1.35 ΔCTCF map). Subtraction of the E1.35 ΔCTCF contact frequency map compared to wildtype CNCCs emphasized these differences, with a large reduction in contact frequency observed along the E1.35 stripe and a general reduction in contact frequency across the TAD (**Figure 4D**; see the arrow on the high confidence map in lower panel emphasizing loss of the E1.35 stripe). From these data, we therefore conclude that E1.35 functions as a structural element that contributes to gene expression, stripe formation and long-range contacts across the locus in a CTCF-site dependent manner. Going forward, we will therefore refer to this element as ‘stripe-associated structural element 1.35’ or SSE1.35 to emphasize the role of this region plays in regulating both domain folding and facilitating target gene expression. Of note, loss of the E1.35 stripe does not lead to ablation of the second TAD-spanning stripe which is present at the barcode overlapping the *SOX9* promoter nor does it disrupt the two TAD boundaries upstream from EC1.45 and downstream from the *SOX9* gene (**Figure 4D**).

To explore the role of the second, promoter-associated stripe at the *SOX9* locus, we again ablated the associated CTCF binding site around 2.2 kb upstream of the *SOX9* TSS using CRISPR/Cas9 genome editing and confirmed loss of CTCF binding (SSEProm ΔCTCF, **Supplementary Figure 4A-B**). Ablation of CTCF binding at this site also had considerable impact on *SOX9* gene expression in CNCCs, with a greater than 20% reduction compared to matched wildtype cells (**Figure 4Bii**). By ORCA imaging, deletion of this CTCF site caused a reduction in TAD-wide contact frequency, though in this case the stripe emanating from the promoter was only weakened, but not fully lost (**Figure 4E**). Subtraction contact frequency maps and bootstrapping confirmed these changes, and further highlighted that the SSE1.35 stripe is retained in the absence of the promoter-associated CTCF site (**Figure 4F**). We therefore named the *SOX9* promoter stripe as ‘stripe-associated structural element at the *SOX9* promoter region’, or SSEProm. Together, we demonstrate that deletion of a CTCF binding site at SSE1.35 or upstream of the *SOX9* promoter (SSEProm) ablates or weakens, respectively, the associated architectural stripe leading to a reduction of contacts across the domain (**Figure 4C-F**). Thus, SSE1.35 and SSEProm function as TAD-internal structural elements that broadly facilitate interactions across the domain.

Ultimately, ablation of either of two structural CTCF sites impacts *SOX9* gene expression in CNCCs. To explore the mechanisms causing a reduction of transcription, we focused on enhancer-promoter proximity for the PRS locus enhancer clusters and the *SOX9* promoter region. Deletion of either CTCF site caused a significant increase in SSE1.35-promoter distances, measured by ORCA imaging (**Supplementary Figure 4C**, p< 0.001, K-S test). By contrast, only promoter-associated CTCF site deletion (at SSEProm), but not SSE1.35 CTCF deletion caused a significant shift in the EC1.45-promoter and EC1.25-promoter pairwise distances (**Supplementary Figure 4C**, p< 0.01, K-S test). However, a general trend towards a reduction in pairwise contact frequency was observed for both CTCF deletions (**Supplementary Figure 4C**, insets). Furthermore, the multiway interactions of the *SOX9* promoter to the three PRS locus regulatory elements were reduced in both SSE1.35 ΔCTCF and SSEProm ΔCTCF lines (**Figure 4G**, additional biological replicates shown in **Supplementary Figure 4D**). Together, these observations are consistent with a model whereby the CTCF-bound SSEs contribute to gene expression by facilitating domain-wide contacts, and as a consequence, promote multiway interactions between the PRS locus elements and the *SOX9* promoter.

One possible explanation for how the SSE1.35 and SSEProm facilitate domain-wide contacts is that they are situated centrally within the 3D domain structure and may promote the compaction of *SOX9* TAD. The average structures derived from our ORCA data in CNCCs (see for example, **Supplementary Figure 1E**) seem to support a central location of the two structural elements. The absolute distance measurements that can be quantified using single chromatin fiber ORCA imaging allows us to test this hypothesis. Indeed, loss of CTCF binding is associated with a significant increase in median radius of gyration (p < 0.001, Wilcoxon rank sum test) for both SSE1.35 ΔCTCF (6.5%) and SSEProm ΔCTCF (7.2%) (**Figure 4H**), indicating a decompaction of the *SOX9* locus in both ΔCTCF lines. To investigate the 3D location of SSE1.35 and SSEProm within the TAD, we plotted the median distance to the geometrical center of the TAD for all imaged regions across the domain (**Figure 4I**). We observed that SSE1.35 and SSEProm are the closest points to the geometrical TAD center, and that deletion of the CTCF binding site at SSE1.35 causes SSE1.35 to move away from the geometrical TAD center (**Figure 4I**). The central positioning of the two SSEs suggests that both SSEs are close to all other regions within the *SOX9* TAD. Indeed, plotting the mean pairwise distance for each imaged region to all other points in the *SOX9* TAD revealed that the shortest mean pairwise distance was for SSE1.35 and SSEProm (**Figure 4J**). Moreover, upon SSE1.35 CTCF deletion, all mean pairwise distances across the domain are increased, further supporting the idea that this structural element promotes the compaction of the TAD (**Figure 4J**). Interestingly, while deletion of the SSEProm CTCF site leads to a similar TAD decompaction and increase in mean pairwise interactions across the TAD (**Figure 4H and 4J**), the *SOX9* promoter remains both the closest point to the geometrical TAD center and to all other regions within the TAD even in the absence of the SSEProm CTCF site (**Figure 4I and J**). This suggests that in contrast to SSE1.35, CTCF binding is not the only feature that promotes the central positioning of the *SOX9* promoter. Together, these data reveal that SSE1.35 and the *SOX9* promoter (SSEProm) are situated centrally within the 3D domain structure, facilitate domain-wide interactions, and promote the formation of a compact structure across the TAD.

### A ‘multi-loop’ model better explains topological features of the *SOX9* locus than a ‘reel-in’ model

To gain mechanistic insight into the processes underlying the central positioning and architectural stripe formation for the two *SOX9* locus structural elements we turned to polymer simulations, which have provided important insights into chromosome topology in previous studies (**Conte et al., 2022**; **Fudenberg et al., 2016, 2017**; **Sanborn et al., 2015**). In particular, based on Hi-C, ChIP and polymer simulations, it has been proposed that architectural stripes that are observed in population level heatmaps are an ensemble feature of the dynamic process of loop extrusion by cohesin via a so-called ‘reel-in’ model (**Vian et al., 2018**). In this model, cohesin is loaded at active enhancers or promoters that are near one or more CTCF sites (**Hsieh et al., 2020**; **Kraft et al., 2019**; **Valton et al., 2021**; **Vian et al., 2018**). Soon after loading, CTCF arrests a single extrusion subunit, while its partner continues to ‘reel-in’ DNA in the opposite direction (**Figure 5A**). In this configuration, cohesin extrudes the intervening DNA in a continuously growing single loop until a second extrusion barrier is encountered. However, from our ensemble analysis, the average CNCC conformation doesn’t resemble a single loop configuration (**Supplementary Figure 1E**), motivating further investigation of the mechanisms driving the observed topological features of the *SOX9* domain.

**Figure 5.**
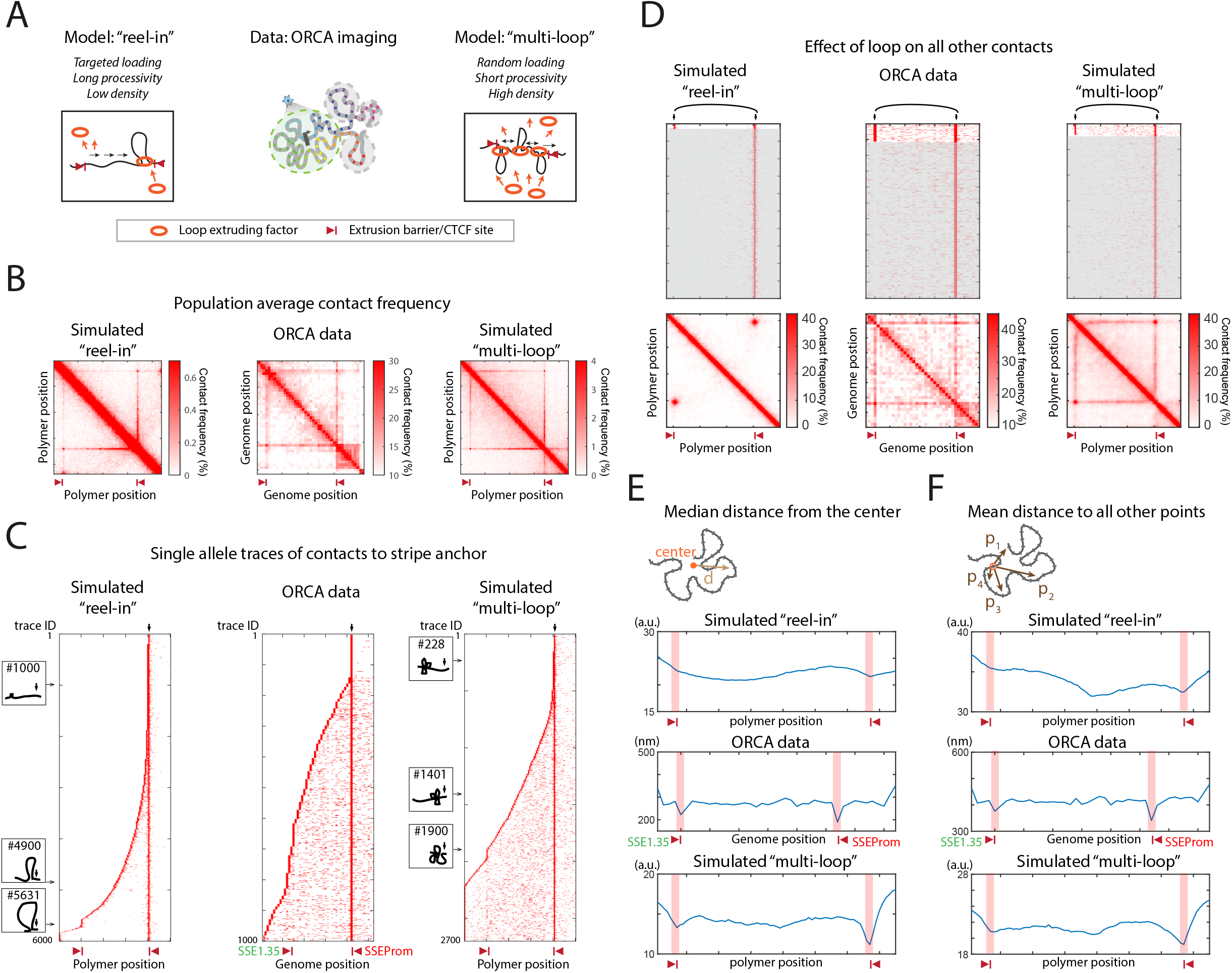
A multi-loop model best fits the observed stripes and central positioning of SSEs at the SOX9 locus. A) Schematics of loop extrusion-based models to explain *SOX9* locus topology. Left, “reel-in” model: cohesin loading is targeted at one site near a loop extrusion barrier with low density of loading and a long processivity. Right, “multi-loop” model: cohesin loading occurs randomly between two loop extrusion barriers at high density with short processivity. B) Population-level contact frequency maps from polymer simulations of the “reel-in’’ model (left) and “multi-loop” model (right), compared to ORCA data from CNCCs (middle, data from Figure 1). C) Single traces of contacts to the right side stripe anchor/loop extrusion barrier. Exemplar traces (#1000, #4900, #5631 for “reel-in” and #228, #1401, #1900 for “multi-loop”) are shown in supplementary Figures 5A and 5B. Cartoons depict a simplified image of the loops for each highlighted trace, with an arrow indicating the viewpoint which is located at the right-hand side stripe anchor. D) Heatmap of traces for pairwise contacts from one stripe anchor across the domain ordered by interactions with a specific point (black arrows) in the domain (upper). Population-level contact frequency maps from traces showing contacts in the upper traces (lower). E) Median distance from the center of the TAD/domain to each point across the domain, averaged across all traces and plotted for each position across the locus. F) The mean distance from a given point to all other points across the domain is plotted for each position across the locus. Red triangle: stripe anchor/loop extrusion barrier.

Our ORCA imaging dataset at the *SOX9* locus provides a unique opportunity to build and compare polymer models to experimental data at both ensemble and single chromatin fiber level. To this end, we built and tested two models using the ‘open2c/polychrom’ framework (**Imakaev et al., 2019**). In the first model, we define that a loop extruding factor (LEF/cohesin) is loaded at a low density, with loading occurring at a single site adjacent to a strong loop barrier, and we allow cohesin to remain bound to DNA and extrude for an extended period, i.e. high processivity. In this configuration, the polymer forms a single loop for each iteration of LEF/cohesin binding, representing loop extrusion-mediated locus scanning as was previously proposed by the ‘reel-in’ model (**Vian et al., 2018**) (**Figure 5A**, left and **Supplementary Figure 5A**). We also considered and simulated a second model, which we call the “multi-loop” model. In this model, to simulate a multi-loop configuration, LEF/cohesin is loaded randomly at any location between the two loop barriers, cohesin loading density is high, yet it only binds and extrudes for a short period, i.e. processivity is relatively low (**Figure 5A**, right and **Supplementary Figure 5B**). When two extrusion barriers are included across a simulated region, both simulations - utilizing the reel-in model or multi-loop model - result in the formation of two stripes in ensemble level maps. These population-level heatmaps are reminiscent of the structures we observed at the *SOX9* locus by ORCA imaging, with two architectural stripes at SSE1.35 and the *SOX9* promoter (SSEProm; **Figure 5B**).

We next explored predictions of these two models at a single chromatin fiber level as the ‘reel-in’ and ‘multi-loop’ models imply different molecular dynamics during the loop extrusion process. The ‘reel-in’ model suggests a dynamic process of a single cohesin complex moving across the domain. To explore the implications of this mechanism, we looked at interactions from one stripe anchor to all points across the domain, sorting the ORCA traces based on the most distal interaction from the stripe anchor. For the simulated reel-in model, the stripe anchor indeed only contacts one point in the domain in most cases (as seen by the white space between the right anchor and the extruding loop, **Figure 5C**, left). To observe how cohesin travels across the TAD to create the observed stripe, we then looked at the ensemble maps from combined single traces that show a contact between the stripe anchor and an arbitrary point. A ‘dot’ like contact is observed in the ensemble map for the reel-in model simulation, revealing that a reeling motion of a single cohesin complex across the TAD creates the stripe observed in the population average (**Figure 5D**, left and **Supplementary Figure 5D)**. By contrast, from our ORCA data in CNCCs we observed multiple intervening contact points from single traces even at a strict threshold for ‘contact’ (<150 nm) (**Figure 5C**, middle and **Supplementary Figure 5C**) and a domain-spanning stripe from ensemble measures (**Figure 5D**, middle). This was in better agreement with the simulated multi-loop model where multiple contact points and a population-level stripe was observed (**Figure 5C**, right and **5D**, right). Together, the multi-loop model is more consistent with our experimental results obtained by ORCA.

Finally, we investigated if the central position of the two structural elements within the 3D domain structure can be recapitulated by either of the two polymer models. From our simulations, we again calculated the predicted distribution of distances from the 3D geometric center of the region to all other points across the domain, as well as the distribution of mean pairwise distances between points across the domain (**Figure 5E**). Similar to our observations from the experimental data, in the multi-loop model simulations, we observed that the loop barriers show the shortest distance to the center of the domain and to the other points within the domain (**Figure 5E and F**, compare middle and lower panels). Indeed, in the multi-loop model simulations, multiple LEFs/cohesins load onto the polymer, stall at loop barriers and consequently results in the loop barriers being frequently positioned in the center of the domain (as seen for a representative simulation, **Supplementary Figure 5E**). This is a unique feature of the multi-loop model, and not observed in the reel-in model simulations (**Figure 5E** and **F**, upper). In summary, our experimental results are better explained by the multi-loop model, suggesting that at large regulatory domains, such as the *SOX9* locus, extreme long-range interactions are facilitated both by the presence of TAD-internal structural elements and by the distributed loading of multiple cohesin molecules. As such, the *SOX9* promoter and its distal enhancers are rarely contained within the single cohesin mediated loop, but rather are joined by a bridge of multiple cohesin molecules stalled against one-another.

## Discussion

Here, we traced 3D locus topology of the *SOX9* regulatory domain for hundreds of cells during differentiation of hESCs to craniofacial progenitors, CNCCs. While overall TAD boundary positions remained constant, we revealed significant changes in the local internal TAD structure during differentiation (**Figure 1**). This observation is in agreement with Hi-C based studies that have demonstrated that while overall TAD structure is typically well-preserved across cell-types (**Dixon et al., 2012**; **Nora et al., 2012**), cell-type specific conformational changes can be observed within TADs (**Barrington et al., 2019**; **Dixon et al., 2015**; **Kraft et al., 2019**; **Phillips-Cremins et al., 2013**; **Smith et al., 2016**). The single-cell, single-chromatin fiber nature of our ORCA locus-tracing datasets enabled us to probe deeper into the variation in locus architecture within and between cell-types. Despite the clearly divergent ensemble structures observed for hESC or CNCC populations of cells, these cell-types could not be distinguished based on the structure alone (**Figure 2**). To further characterize *SOX9* domain conformational heterogeneity during the hESC to CNCC differentiation, we performed clustering of single chromatin fiber traces from these two cell-types, and observed that all resultant structural clusters were in fact represented by both hESCs and CNCCs. However, certain conformations were more common in one or the other cell-type, suggesting that ensemble-average differences in locus topology arise through cell-type specific changes in frequency of commonly sampled conformations (see **Figure 6A** for model schematic). Together, we highlight the flexible, likely dynamic nature of locus topology across a population of cells, which is perhaps influenced by distinct cell-type specific epigenetic and transcriptional states.

**Figure 6.**
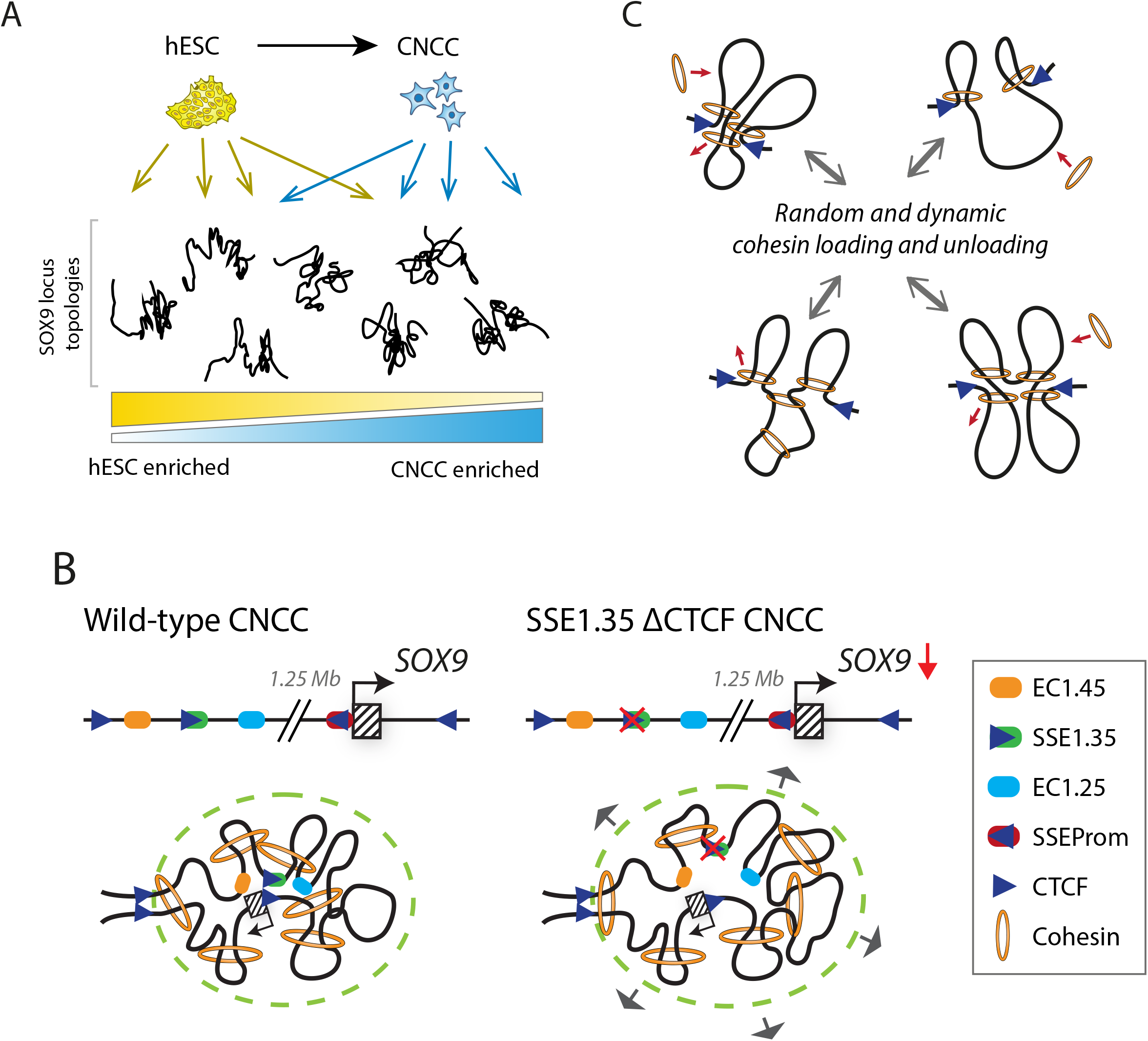
Schematic models of dynamic domain topology during CNCC differentiation, and the role of structural elements and multi-loop formation in SOX9 locus topology. A) Single trace analysis of ORCA data from hESCs and CNCCs revealed that ensemble-average differences in locus topology arise through cell-type specific changes in frequency of conformations commonly sampled by both cell-types. B) In wildtype CNCCs, the *SOX9* promoter sits close to the middle of the TAD. SSEProm and SSE1.35 element are bound by CTCF and interact frequently drawing SSE1.35 into the center of the domain. Multiple cohesin loading events act to compact the domain. Ablation of the CTCF binding sites at SSEs leads to reduction of central SSE positioning within the TAD, loss or reduction of associated architectural stripes, increase in domain radius and reduction in *SOX9* expression. C) ORCA imaging data and simulations support a ‘multi-loop’ model with two important parameters, firstly cohesin is loaded randomly without bias between the two loop barriers and secondly multiple cohesin complexes can extrude chromatin within the domain simultaneously.

One of the most striking features of the ensemble interaction frequency maps for the *SOX9* TAD was the appearance of two TAD-spanning stripes in CNCCs. We identified two stripe-associated structural elements (SSEs), one at the PRS locus element SSE1.35 (1.35 Mb upstream of *SOX9*) and the other upstream and proximal to the *SOX9* promoter (SSEProm). These SSEs promote the formation of the observed stripes, are bound by CTCF and facilitate expression from the *SOX9* gene. Therefore, we define a new regulatory function for SSE1.35, that we had previously shown is neither a canonical enhancer (despite its enrichment for enhancer-associated chromatin features) (**Long et al., 2020**) nor an enhancer booster (**Figure 3**), and further identify a second structural element proximal to the *SOX9* gene. There are a number of parallels between the two SSEs identified here and tethering elements recently reported in Drosophila development (**Batut et al., 2022**). Both SSEs and Drosophila tethering elements are associated with open chromatin, do not overlap with TAD boundaries, and impact gene expression when ablated despite not having functional enhancer activity. However, there are also many notable distinctions, including that the *SOX9* locus SSEs are CTCF-dependent and associated with stripes rather than points/dots on contact frequency maps (**Figure 4**). This suggests that the Drosophila tethering elements may form relatively frequent loops between two specific elements, while the *SOX9* locus SSEs described here facilitate broad domain-wide interactions.

Promoter-proximal CTCF binding has also been demonstrated in mouse ESCs and NPCs, and human cancer cell lines to promote distal enhancer-dependent gene activation for many genes by facilitating contact of promoter and distal enhancers that lie between CTCF sites (**Kubo et al., 2021**; **Schuijers et al., 2018**). In contrast, at the *SOX9* locus two CTCF-bound SSEs sit in between (rather than flanking) the strongest distal enhancer (EC1.45) and its target gene, therefore the *SOX9* locus SSEs appear to mediate enhancer activity from both upstream and downstream of their long-range interaction. Notably, this is also distinct from regulatory neighborhoods which appear to act through an insulation-based mechanism that brings enhancers and promoters into a shared domain (**Dowen et al., 2014**). We propose that instead of promoting insulation or directionality, the *SOX9* locus SSEs facilitate function of distal ECs in *SOX9* gene activation through enabling the formation of a multi-loop structure and compaction of the entire domain (see **Figure 6B** for model schematic).

We observe a hub-like conformation at the *SOX9* locus which features a high frequency of ORCA traces with the *SOX9* gene located close to the center of the domain. By virtue of its central positioning, the promoter appears to be in an optimal location to sample interactions with all other regions across the domain. The CTCF site at SSE1.35 also draws the distal region of the TAD into the center of the domain. The two SSEs are not entirely equivalent however, as the central positioning of SSE1.35 appears less pronounced than for SSEProm, and both the central positioning and stripe are lost for SSE1.35 upon CTCF site deletion. By contrast, the promoter-adjacent stripe is less dependent on CTCF binding, suggesting partial redundancy with other mechanisms mediating its formation. For example, RNA Pol II has also been shown to block cohesin movement on chromatin, thus serving as an additional extrusion barrier at transcribing genes (**Banigan et al., 2022**; **Busslinger et al., 2017**), and emerging evidence suggests that extrusion might also be obstructed by other complexes such as the replication machinery (**Dequeker et al., 2022**; **Jeppsson et al., 2022**). Another possibility is that the central positioning of the *SOX9* gene within the TAD is additionally facilitated via a loop extrusion-independent mechanism, such as association with transcription factories (or other nuclear bodies) or by polymer phase separation, as proposed by the Strings and Binders (SBS) model (**Conte et al., 2022**).

To explore how architectural stripes and regulatory hubs may form, we used polymer simulations to compare our empirical ORCA data to an existing ‘reel-in’ model that was proposed based on population level Hi-C heatmaps, and emphasized targeted loading of cohesin (**Kraft et al., 2019**; **Valton et al., 2021**; **Vian et al., 2018**). Counter to the ‘reel-in’ model predictions, we observed multiple interactions across the *SOX9* domain between SSE1.35 and the *SOX9* promoter. Our ORCA results and simulations instead support a ‘multi-loop’ model with two important parameters, firstly that cohesin complexes are loaded randomly without bias between the two loop barriers and secondly that multiple cohesin complexes can extrude chromatin within the domain simultaneously. This agreement implies that loop-extruding cohesin complexes are dense enough and processive enough at the *SOX9* locus to collide frequently and that when they collide, they do not proceed past one-another (though they need not be processive enough to walk the intervening 1.35 Mb distance without falling off). If there were not collisions, the CTCF anchored SSE1.35 and *SOX9* promoter would not be enriched in multiple contacts to the intervening domain.

Evidence for the preferential loading of cohesin at promoters or active enhancers comes from ChIP-seq for the proposed cohesin loader, NIPBL (**Fudenberg et al., 2016**; **Gullerova & Proudfoot, 2008**; **Kagey et al., 2010**; **Kraft et al., 2019**; **Vian et al., 2018**). However, antibody specificity for endogenous NIPBL protein has been questioned, and a recent study that endogenously-tagged NIPBL has instead indicated a more uniform loading of cohesin (**Banigan et al., 2022**), in keeping with our first proposed parameter of the multi-loop model. Of note, NIPBL has been reported to have roles in addition to cohesin loading, and so future work is required to more carefully map sites of cohesin loading (**Bauer et al., 2021**; **Davidson & Peters, 2021**).

Regardless, polymer simulations using uniform cohesin loading more closely reproduced the experimental Hi-C maps than preferential loading of cohesin at the promoter (**Banigan et al., 2022**). Furthermore, it has recently been estimated from Micro-C data that cohesin processivity is around 150 kb (**Gabriele et al., 2022**; **Gassler et al., 2017**). This processivity would not be sufficient for a cohesin complex to extrude across the entire 1.25 Mb between the *SOX9* gene and distal ECs if cohesin only loaded at promoters or enhancers as proposed in the ‘reel-in’ model. Reported measures of cohesin processivity therefore support the second requirement of the multi-loop model for multiple simultaneous cohesin extrusion events. Lastly, the ‘multi-loop’ model not only best fits our empirical data at the *SOX9* locus, it also predicts the central positioning of the two stripe-associated structural elements within the regulatory domain (**Figure 5**). Together, our proposed ‘multi-loop’ model best recapitulates our experimental ORCA data by multiple measures and is supported by recent biophysical and genomic measures of cohesin processivity and loading.

Overall, our ORCA data and polymer simulations suggest that the stripe pattern observed at the population level is not an ensemble feature of the dynamic process of a single extruding loop (‘reel-in’). Instead, the stripe appears to be built from multiple loops across the domain that also act to frequently tether the SSE1.35 element to the *SOX9* promoter via multiple cohesin bridges (see **Figure 6C** for model schematic). The multiple loop configuration provides several advantages over the ‘reel-in’ model, especially in extreme long-range gene regulation such as at the *SOX9* locus. As described, the ‘multi-loop’ model overcomes the low reported processivity of cohesin by the distributed cohesin loading and multiple cohesin mediated loops that together compact the domain and bring the promoter and enhancers into proximity. Moreover, multiple cohesin complexes extruding in the domain imply faster and better efficiency in mediating promoter-enhancer interaction compared to a single extruding factor scanning along the domain in the ‘reel-in’ model. Multiple cohesin configuration also buffers the effect of stochastic loading/unloading of cohesin, providing robust interactions. Frequent positioning at the center of the domain allows *SOX9* gene proximity to the rest of the domain including all enhancers. Together, the ‘multi-loop’ model provides a conceptual framework for understanding how structural elements can become centrally situated in a regulatory domain to enable domain-wide interactions, which in turn may facilitate robust regulation of gene expression, especially at large TADs with extremely distal enhancers.

Craniofacial development appears to be highly sensitized to alterations in levels of *SOX9* expression in CNCCs, given the selectivity of phenotypes in patients with *SOX9* haploinsufficiency and the severe developmental consequences of loss of non-coding DNA and distal regulatory enhancers at the PRS locus (**Benko et al., 2009**; **Long et al., 2020**). In our *in vitro* model of CNCC development, a 20% reduction in *SOX9* expression was observed upon loss of either SSE1.35 or SSEProm, a level of dysregulation that is sufficient to drive phenotypic change in mouse models (**Long et al., 2020**). Therefore, as more genome sequences will become available from patients with PRS and related dysmorphologies in the future, we postulate it might be worthwhile to scan for mutations affecting CTCF binding at the two SSEs identified in this study.

## Acknowledgements

We thank members of the Wysocka lab and the Boettiger lab for comments and discussion. We thank Dr. Raquel Fueyo, Dr. Sahin Naqvi, Dr. Antonina Hafner, Sedona Murphy, and Kirsty Uttley for critical reading of the manuscript. This work was supported by HHMI (to J.W.), R35 GM131757 (to J.W.), U01 DK127419 (to J.W. and A.N.B.), Lorry Lokey endowed professorship (to J.W.), Beckman Young Investigator Award (to A.N.B.), Packard Fellowship for Science and Engineering (to A.N.B.), Wellcome Trust Sir Henry Wellcome Fellowship 106051/Z/14/Z (to H.K.L.) and Damon Runyon Postdoctoral Research Fellowship (to M.P.).

## Author Contributions

Conceptualization, L.F.C., H.K.L., A.N.B. and J.W.; Methodology, L.F.C., H.K.L, A.N.B., and J.W.; Formal Analysis, L.F.C., H.K.L., T.S., and A.N.B.; Investigation, L.F.C., H.K.L., and M.P.; Resources, L.F.C. and H.K.L.; Writing – Original Draft, L.F.C. and H.K.L.; Writing – Review & Editing, all authors; Visualization, L.F.C., H.K.L., and A.N.B.; Supervision, A.N.B. and J.W.; Funding Acquisition, H.K.L., M.P., A.N.B., and J.W.

## Data and code availability

All probe coordinates and data tables are in the process of being annotated and formatted for upload on the 4DN public access server. Code for image analysis and polymer simulations will be available at https://github.com/BoettigerLab/sox9-ORCA-2022. Polymer simulations require the polychrom simulation tools, available at: https://github.com/BoettigerLab/polychrom. Additional information on polymer simulation modelling is included in the Methods.

## Methods

### EXPERIMENTAL MODEL AND SUBJECT DETAILS

#### Culturing human embryonic stem cells (hESCs)

Human embryonic stem cells (hESCs) were purchased from ATCC (WA09, also known as H9; RRID: CVCL_9773). hESCs were fed every day with mTeSR (Stem Cell Technologies), grown on Matrigel Growth Factor Reduced (GFR) Basement Membrane Matrix (Corning) at 37°C and passaged every 5-6 days using ReLeSR (Stem Cell Technologies).

### METHOD DETAILS

#### Genome-editing of hESC cell lines using CRISPR/Cas9

Two edited hESC cell lines were generated previously, ΔEC1.45/+ and ΔEC1.25/+ (**Long et al., 2020**). Targeted deletion of E1.35 was performed in a similar manner. Briefly, H9 hESCs were transfected using FuGENE 6 (Promega) with a targeting construct containing Blasticidin selection cassette, flanked by FRT sites, and homology arms for either side of E1.35 along with a plasmid encoding Cas9 plus single guide RNAs (sgRNAs) targeting sites flanking E1.35. Transfected hESCs were grown to confluency and split onto a new plate before selection with 1 mg/mL Blasticidin until all cells died on a mock/GFP transfected control well. Surviving colonies were picked into a 48-well plate, expanded, passaged and screened for enhancer deletion using a genomic primer and a primer within the targeting cassette. Heterozygous enhancer deleted clones were transfected with a Flippase-expressing plasmid and clones screened for excision of the selection cassette by PCR. For screening, genomic DNA was extracted using QE buffer (Lucigen) and PCR was performed using Q5 polymerase (NEB). Heterozygous enhancer deletions were generated to facilitate allele-specific *SOX9* gene expression analysis.

#### Differentiation of hESC to CNCCs and chondrocytes

hESCs were differentiated to human cranial neural crest cells (CNCCs) as described previously (**Long et al., 2020**; **Prescott et al., 2015**). Briefly, hESCs were grown for 5-6 days until large colonies formed, which were disaggregated using collagenase IV and gentle pipetting. Clumps of ∼200 hESCs were washed in PBS and transferred to a 10cm Petri dish in neural crest differentiation media (NDM). NDM: 1:1 ratio of DMEM-F12 and Neurobasal, 0.5x Gem21 NeuroPlex Supplement With Vitamin A (Gemini, 400-160), 0.5x N2 NeuroPlex Supplement (Gemini, 400-163), 1x antibiotic/antimycotic, 0.5x Glutamax, 20ng/ml bFGF (PeproTech, 100-18B), 20ng/ml EGF (PeproTech, AF-100-15) and 5ug/ml bovine insulin (Gemini Bio-Products, 700-112P). After 7-8 days, cranial neural crest cells (CNCCs) emerged from neural spheres that had attached to the Petri dish. After 11 days, CNCCs were passaged onto fibronectin-coated 6-well plates using accutase and fed with neural crest maintenance media (NMM). NMM: 1:1 ratio of DMEM-F12 and neurobasal, 0.5x Gem21 NeuroPlex Supplement with Vitamin A (Gemini, 400-160), 0.5x N2 NeuroPlex Supplement (Gemini, 400-163), 1x antibiotic/antimycotic, 0.5x Glutamax, 20ng/ml bFGF, 20ng/ml bFGF EGF and 1mg/ml BSA (Gemini, 700-104P). After 2-3 days, CNCCs were split 1:3 and the following day cells were fed with neural crest long-term media. Long term media: neural crest maintenance media plus 50pg/ml BMP2 (PeproTech, 120-02) and 3uM CHIR-99021 (Selleck Chemicals, S2924). CNCCs were then passaged twice to passage 4 and processed for various assays.

#### cDNA preparation and reverse transcriptase digital droplet PCR (RT-ddPCR)

Total RNA was extracted from late (passage 4, P4) CNCCs differentiated from hESC wild-type or mutant cell-lines using Trizol reagent (Invitrogen) for at least two differentiations. 500ng – 1ug RNA was used to generate cDNA using the SuperScript Vilo IV MasterMix with ezDNase enzyme (Invitrogen, 11766050). Primers and locked nucleic acid (LNA) probes were designed by the IDT custom design service to the human SOX9 3’UTR, centered on the rs74999341 T/C SNP – a HEX LNA probe detects the T-allele and a FAM LNA probe detects the C-allele. cDNA dilution factor was determined using qPCR with 1X PrimeTime Gene Expression Master Mix (IDT), 500 nM primers and 250 nM probes, run on LightCycler 480 (Roche). ddPCR reactions were performed using diluted cDNA (10-100X diluted), 900 nM primers and 250 nM probes and 1X ddPCR Supermix for probes (no dUTP, BioRad). ddPCR droplets were generated using the QX200 Droplet Generator (BioRad) and droplets were read using QX200 Droplet Reader (BioRad) and analyzed using the QuantaSoft Software (BioRad).

#### Chromatin immunoprecipitation digital droplet PCR (ChIP-ddPCR) or ChIP-qPCR

5-15 million cells were cross-linked per ChIP experiment in 2mL PBS per 6-well with 1% methanol-free formaldehyde for 5-10 min and quenched with a final concentration of 0.125M glycine for 5 min with nutation. Cross-linked cells were washed with PBS, scraped and pelleted by centrifugation, flash-frozen in liquid nitrogen and stored at -80°C. Samples were defrosted on ice and resuspended in 5mL LB1 (50 mM HEPES-KOH pH 7.5, 140 mM NaCl, 1 mM EDTA, 10% glycerol, 0.5% NP-40, 0.25% Triton X-100, with 1X cOmplete Protease Inhibitor Cocktail and 1 mM PMSF) and rotated vertically for 10 min at 4°C. Samples were centrifuged for 5 min at 1350 x g at 4°C, and resuspended in 5mL LB2 (10 mM Tris, 200 mM NaCl, 1 mM EDTA, 0.5 mM EGTA, with 1X cOmplete Protease Inhibitor Cocktail and 1mM PMSF) and rotated vertically for 10 min at 4°C. Samples were centrifuged for 5 min at 1350 x g at 4°C, and resuspended in 1mL LB3 per 10 million cells (up to 1 mL per ChIP). Samples were sonicated in 1mL AFA tubes for 5 min on E220 evolution Covaris with settings Peak power = 140, Duty Factor = 10, Cycles per burst = 200 to achieve chromatin sized approximately 500-2000bp. Following sonication, samples were recombined (if aliquoted for sonication), Triton X-100 was added to the fragmented chromatin to a final concentration of 1%, and the chromatin divided for input (1%–2%) and ChIP samples. 5 mg anti-histone H3K27ac (Active Motif, 39133) antibody or 10 μL anti-CTCF (Cell Signaling, 2899S) antibody was added per ChIP sample, and incubated overnight at 4°C with rotation. Protein G Dynabeads (ThermoFisher) were first blocked with Block solution (0.5% BSA (w/v) in 1X PBS) and then added to cleared chromatin to bind antibody-bound chromatin during a 4-6 hour incubation. Chromatin-bound Dynabeads were washed at least 6 times with chilled RIPA wash buffer (50 mM HEPES-KOH pH 7.5, 500 mM LiCl, 1 mM EDTA, 1% NP-40, 0.7% Na-Deoxycholate), followed by a wash with chilled TE + 50 mM NaCl. Chromatin was eluted for 15-30 min in Elution Buffer (50 mM Tris, 10 mM EDTA, 1% SDS) at 65°C with frequent vortexing. The ChIP and input samples were then incubated at 65°C overnight to reverse cross-links (12-16 hours). Samples were diluted and sequentially digested with RNase A (0.2 mg/mL) for 2 hours at 37°C followed by Proteinase K (0.2 mg/mL) for 2 hours at 55°C to digest protein. ChIP and input samples were purified by phenol-chloroform-isoamylalcohol extraction and precipitation with final concentration 70% ethanol, 0.3M NaOAc pH 5.2 and 1.5 mL glycogen.

For ChIP-ddPCR, primers were designed to amplify across the EC1.45, E1.35 and EC1.25 regions, and allele-specific probes were designed to distinguish between wild-type and mutated alleles. ChIP and input dilution factors were determined using qPCR with 1X PrimeTime Gene Expression Master Mix (IDT), 500 nM primers and 250 nM probes, run on LightCycler 480 (Roche). ddPCR reactions were performed using ChIP DNA (40X diluted) and input DNA (640X diluted), 900 nM primers and 250 nM probes and 1X ddPCR Supermix for probes (no dUTP, BioRad). ddPCR droplets were generated using the QX200 Droplet Generator (BioRad) and droplets read using QX200 Droplet Reader (BioRad) and analyzed using the QuantaSoft Software (BioRad).

For ChIP-qPCR, primers were designed to amplify adjacent to the SSE1.35 and SSEProm CTCF site. ChIP DNA and input DNA were diluted 5X and amplified using 500 nM primers and SensiFAST SYBR (No-ROX Kit, Bioline) using a LightCycler 480 (Roche).

#### ORCA

The *SOX9* ORCA probes, with 80 barcodes tiling the surrounding locus (Supplementary Table S1-2), were designed as previously described (**Mateo et al., 2019, 2021**).

#### Sample preparation

ORCA imaging experiments were performed following the protocol described (**Mateo et al., 2019**) and in detail in (**Mateo et al., 2021**). Briefly, hESCs and CNCCs were fixed in 4% PFA in 1x PBS for 10 min. Fixed cells were plated on a poly-D-lysine-coated 40 mm coverglass and permeabilized with 0.5% Triton-X in 1x PBS for 10 min, followed by 2 washes with 1x PBS. Cells were incubated for 5 minutes in 0.1 M HCl, followed by 3 washes with 1x PBS and 3 washes with 2x SSC. We then treated cells for 35 min in 2x SSC + 50% vol/vol formamide and 0.1% Tween. 3 μg SOX9 ORCA probe in 30 μl Hybridization buffer (2x SSC, 50% vol/vol formamide and 0.1% Tween) were added onto cells. Cells were then denatured for 3min at 90°C and incubated overnight at 42°C. After the hybridization, cells were washed twice for 10min in 2x SSC at 42°C, then postfixed in 2% GA, 8% PFA in 1x PBS for 30 min. Cells were then washed in 2x SSC and either imaged directly or stored for up to a week at 4°C.

#### Image acquisition

Samples were imaged on the custom microscopy and microfluidics setup as described in (**Mateo et al., 2019, 2021**). Briefly, samples were mounted into a flow chamber. Readout barcodes were visualized sequentially by complementary oligos carrying Cy5 dye followed strand displacement and washes. A Cy3 oligo that label all barcodes were also imaged at each round of imaging as fiducial spots. The fiducial spots were subsequently used for spot calling and registration in the image analysis pipeline.

#### Image processing

Spot calling, localization, and drift correction were performed as previously described. (**Mateo et al., 2019, 2021**). Software for processing the raw data is available at https://github.com/BoettigerLab/ORCA-public

##### ORCA image analysis

After spot calling and localization, barcodes with low hybridization efficiency (< 10%) were filtered. Traces with low number of barcodes labeled (< 30%) and very high number of barcodes labeled (> 90%) were then filtered. Pairwise distances were calculated for all filtered barcodes and traces. Contact frequency is defined as the percentage of pairwise inter-probe distances below 250 nm. To estimate the confidence interval for the difference in contact frequency between conditions, we performed bootstrapping by randomly sampling with replacement from the data.

To estimate the volume of the chromatin domain, we calculated the Radius of gyration of each trace. **Radius of gyration (Rg)** of the chromatin domain was computed from the vector data of molecule locations as 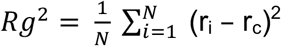

Where r_i_ is a vector of the coordinate of an individual localization and r_c_ is the centroid of all N localizations.

To estimate the average 3D structure of domain, we performed **non-metric multidimensional scaling (MDS)** on the average pairwise distance matrix from ORCA. We calculated Euclidean distances for all pairs of points in 3D using Kruskal’s normalized stress criterion.

##### K-means clustering and tSNE

We used the pairwise distance from the points in *SOX9* TAD to *SOX9* gene (barcodes 11-52 to barcode 42). We only used traces that were labeled by more than 60% barcodes and interpolated the missing data points in these traces using the mean pairwise distance from the neighboring two barcodes. K-means clustering with different number of clusters was performed on the interpolated data and the optimal number of clusters was determined by the Elbow method. Clusters from K-means clustering were superimposed on the tSNE plots.

### Polymer simulation

Polymer simulations were conducted using the open source polychrom software (**Imakaev et al., 2019**) from open2c, which builds on previously published approaches for simulating chromatin structure and loop extrusion (**Fudenberg et al., 2016**; **Nuebler et al., 2018**). This software constructs Langevin dynamic simulations of flexible polymers moving under thermal noise with user defined energy potentials to describe the bending stiffness, bond stiffness, and molecular interactions among monomers, including the links produced by loop extrusion factors. The simulations use the freely distributed openMM framework (**Eastman et al., 2017**), widely used in molecular dynamics and protein structure simulations, to provide GPU accelerated computation. To simulate the “reel-in” model of stripe formation, we added targeted loading to this framework. This expanded version of the software is available at https://github.com/BoettigerLab/polychrom. Annotated, executable scripts with the parameters used to simulate both the “reel-in” and the “multi-loop” models will be provided at https://github.com/BoettigerLab/sox9-ORCA-2022.

Briefly, both the “reel-in” and the “multi-loop” simulations used a polymer chain consisting of 1000 monomers, with extrusion blocking elements (simulating the CTCF structural elements) positioned internally (at monomer 200 and 800). In the “reel-in” model, cohesin was primarily loaded at monomer 205 and 795. To explore the relative difference in loading strength, each cohesin loaded had a 10% probability of loading at monomer 205 and 90% probability of loading at monomer 795. In the “multi-loop” model cohesin loaded randomly throughout the domain instead. In the “reel-in” model, no more than two LEFs/cohesins were extruding at any given time, and LEFs/cohesins remained bound for an average of 800 monomer steps. In the “multi-loop” model up to 20 LEFs/coheins could be randomly loaded across the domain, but remained on the polymer long enough on average to traverse only 40 monomer steps. Capture probability for an extrusion blocking element to stop a LEF/cohesin, average residence time of a LEF/cohesin at the capture site, and polymer mechanical properties were otherwise identical between both simulations. Further details can be seen in deposited scripts.

### QUANTIFICATION AND STATISTICAL ANALYSIS

#### Analyzing and plotting ddPCR data

Concentration of the two alleles of *SOX9* was determined by RT-ddPCR using allele-specific HEX/FAM probes from the QuantaSoft Software. To plot allelic skew, concentration for the mutated allele was divided by the concentration for the wild-type allele and plotted as a ratio (red boxplots). For matched wild-type cells, the same ratio was calculated, and then the values were normalized such that the wildtype median equals to 1.

##### External datasets

External HiC data was obtained from the 4DN consortium: 4DNESRJ8KV4Q. CNCC ChIP-seq datasets are from (**Long et al., 2020**; **Prescott et al., 2015**).

## Abbreviations

(NCC): neural crest cell
(CNCC): cranial neural crest cell
(PRS): Pierre Robin sequence
(TAD): topologically associating domain
(ORCA): optical reconstruction of chromatin architecture
(SSE): stripe-associated structural element

---

**Supplementary Figure 1.**
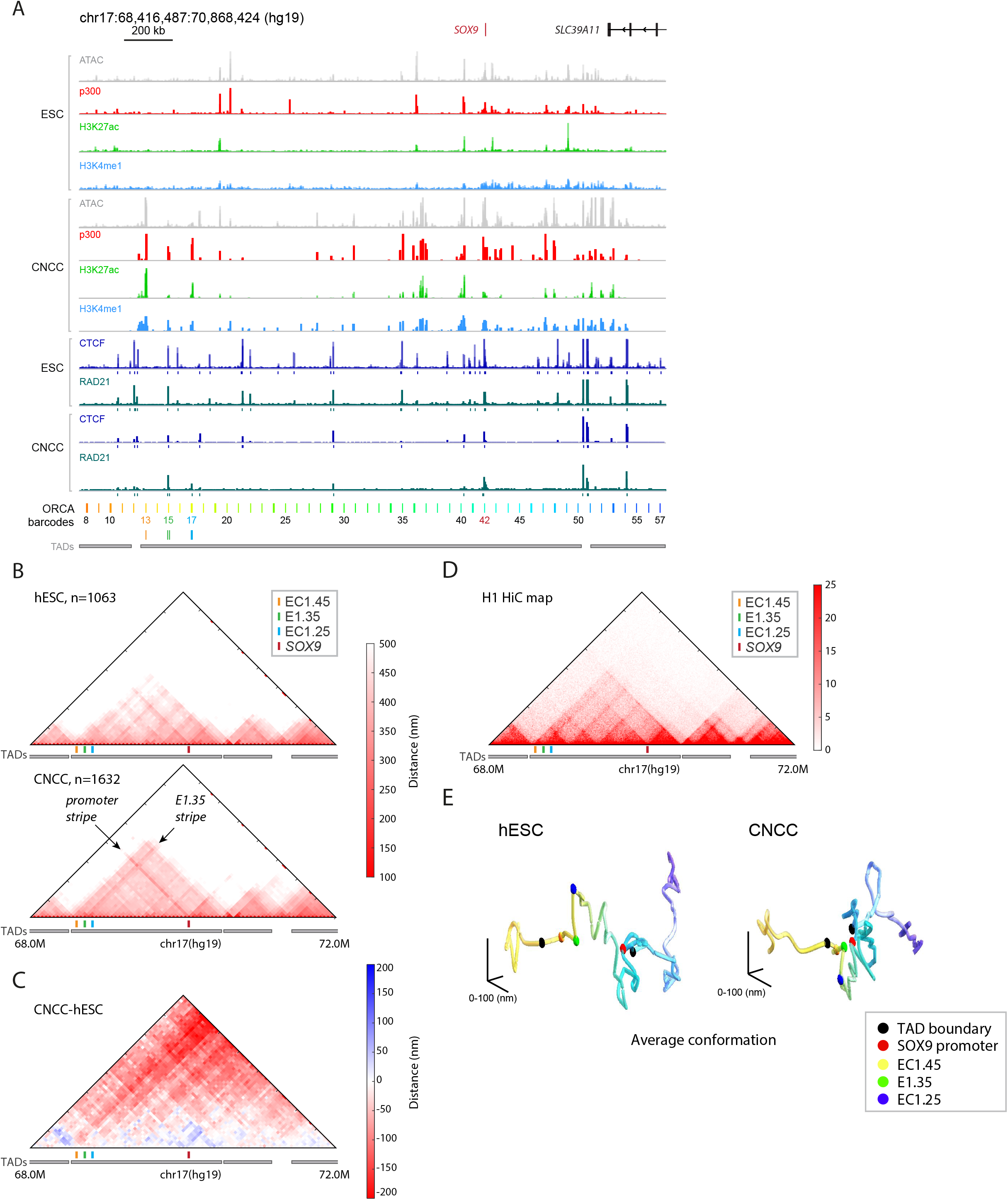
A) ATAC-seq and p300, H3K27ac, H3K4me1, CTCF and cohesin ChIP-seq data from hESCs and CNCCs for the locus surrounding the *SOX9* gene. Enhancer clusters EC1.45 and EC1.25 are highlighted along with the nearby E1.35 element. ORCA barcodes are annotated across the domain, along with TADs. B) Absolute pairwise distance maps for all ORCA barcodes across the *SOX9* locus from ORCA imaging data for hESC (upper) and CNCCs (lower). C) Absolute pairwise distance subtraction maps for CNCC-hESC. D) Hi-C contact frequency maps for H1 hESCs. E) MDS plots of the average 3D structure of the *SOX9* locus and flanking regions in hESCs (left) and CNCCs (right). Barcodes are highlighted that overlap interesting genomic features. Data from Figure 1B.

**Supplementary Figure 2.**
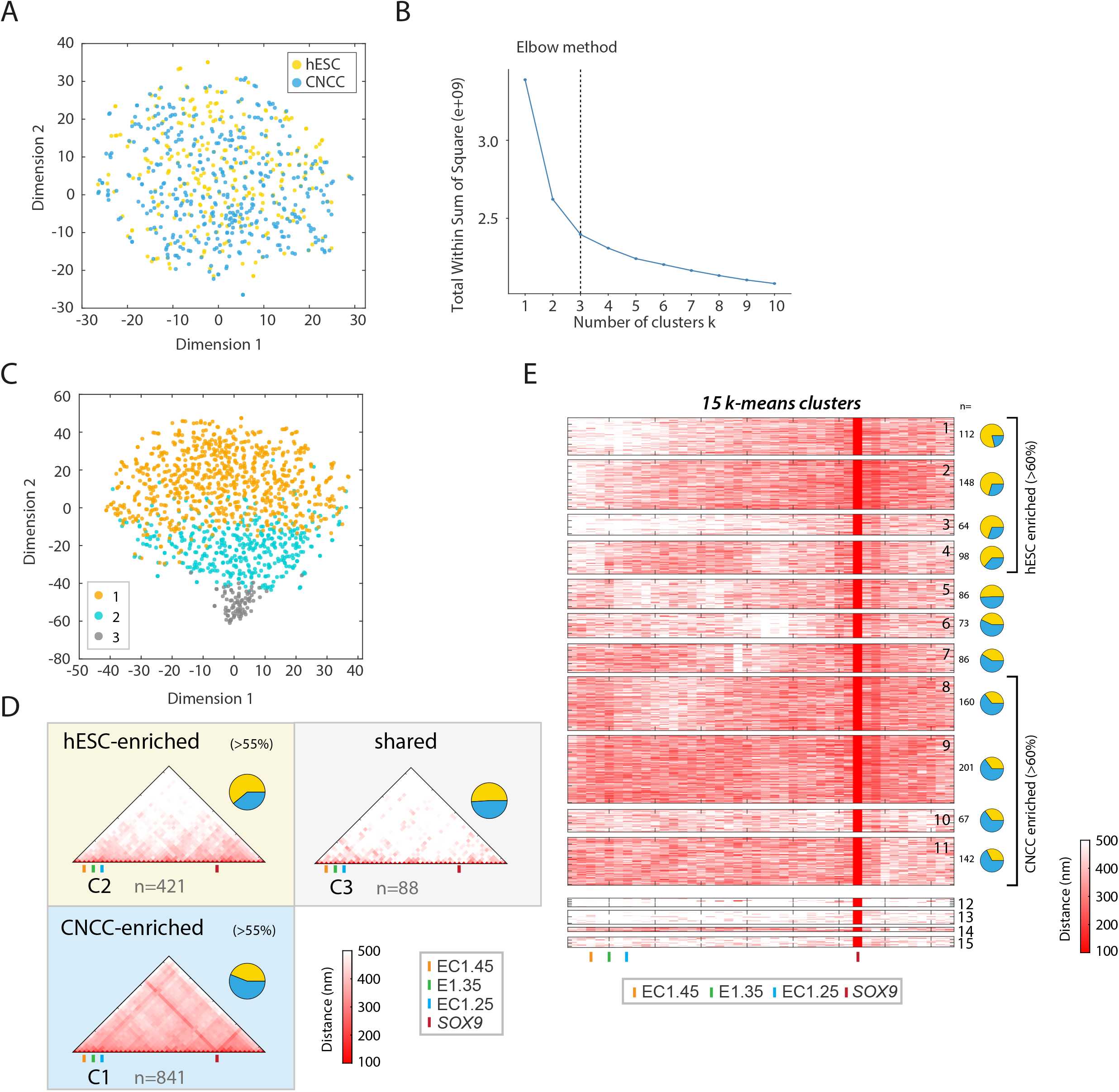
A) A tSNE plot representing all pairwise distances of all versus all positions across the *SOX9* TAD (data from Figure 1). B) Elbow method to determine optimal number of K-means clusters for all pairwise distances of the *SOX9* promoter to the remainder of the TAD (data from Figure 1). C) tSNE plot from Figure 2A with 3 clusters from K-means clustering indicated. D) Contact frequency maps from all traces for each cluster from C. E) Heatmaps representing absolute pairwise distances of the *SOX9* promoter with all other regions in the surrounding domain, showing all traces from each of the 15 K-means clusters from Figure 2B.

**Supplementary Figure 3.**
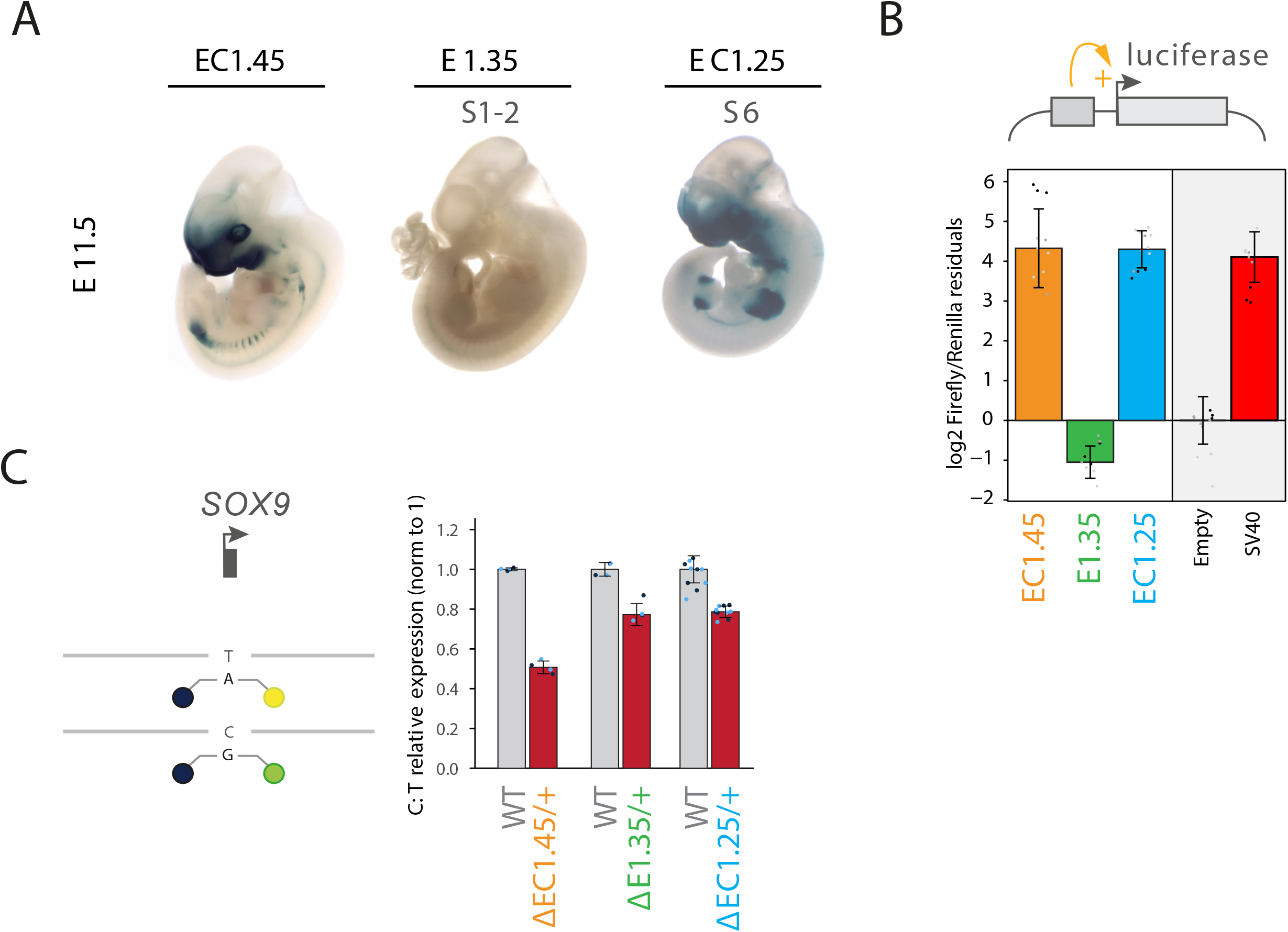
A) LacZ reporter assay for EC1.45, E1.35 and EC1.25 at embryonic day E11.5. B) Luciferase assay for EC1.45, E1.35 and EC1.25 in CNCCs. C) Allelic expression of *SOX9* for wildtype and heterozygous EC1.45, E1.35 and EC1.25 mutant CNCCs, plotting mutant versus wildtype allelic expression measured by RT-ddPCR, with wildtype expression normalized to 1. All data (apart from E1.35 deletion expression data) from (**Long et al., 2020**).

**Supplementary Figure 4.**
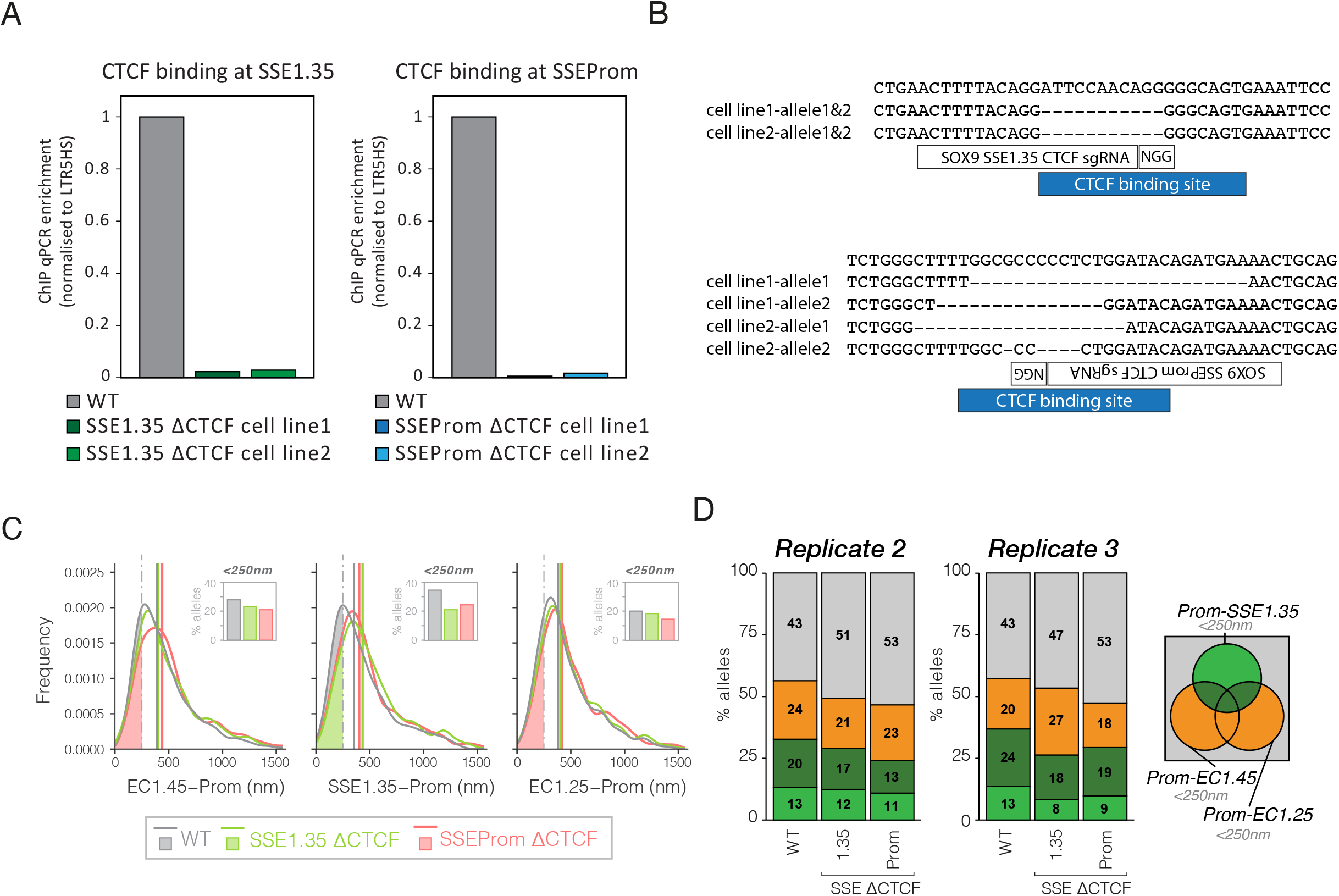
A) ChIP-qPCR for CTCF at SSE1.35 for two homozygous mutant clones compared to wildtype (left), and SSEProm for two homozygous mutant clones compared to wildtype (right). B) Multiple sequence alignment illustrating the indel mutations for each of the alleles for the homozygous CTCF mutant clones. C) Frequency plots showing pairwise distance distributions for wild-type, SSE1.35 ΔCTCF and SSEProm ΔCTCF CNCCs for EC1.45, E1.35 (SSE1.35) and EC1.25 to the *SOX9* promoter from left to right. Percentage of alleles with respective pairwise distances <250 nm are shown as an inset. D) Two additional replicates for Figure 4I. Percentage of alleles with promoter proximity to EC1.45, SSE1.35 and/or EC1.25 including multiway interactions for wild-type, SSE1.35 ΔCTCF and SSEProm CTCF CNCCs.

**Supplementary Figure 5.**
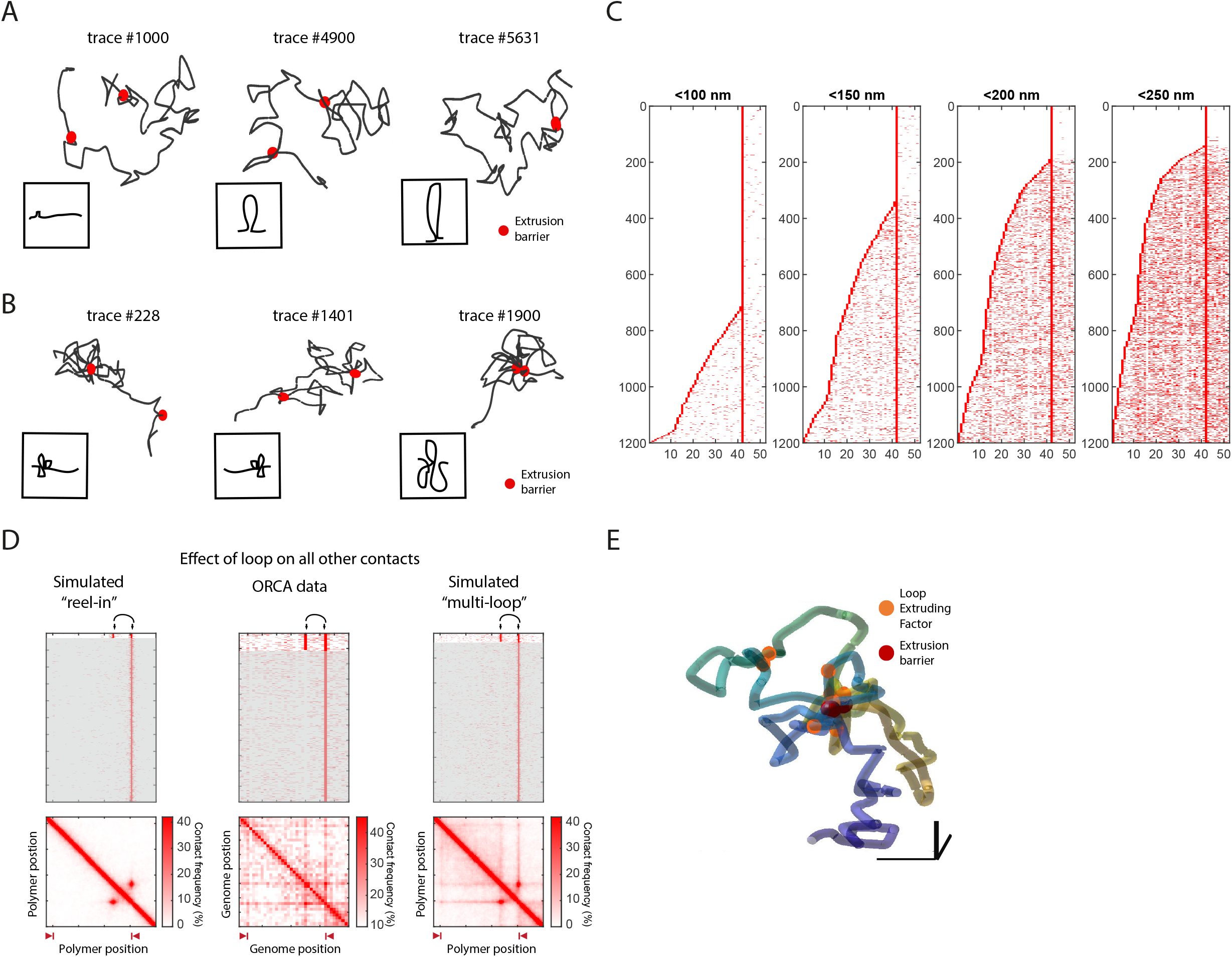
A) Exemplar traces, #1000, #1900 and #5631 from the “reel-in” model. B) Exemplar traces, #228, #1401 and #1900 from the “multi-loop” model. C) Single chromatin ORCA traces of contacts to the stripe anchor at the right side of the domain at different threshold for contacts as indicated above each plot. D) As Figure 5D. Heatmap of iterations (polymer simulations) or traces (ORCA imaging) for pairwise contacts from one stripe anchor across the domain ordered by interactions with a specific point (black arrows) in the domain (upper). Population-level contact frequency maps from traces showing contacts in the upper traces (lower). E) An exemplar trace from the “multi-loop” model showing multiple loop conformation and central positioning of the stripe anchors.

